# Structure and functionality of a multimeric human COQ7:COQ9 complex

**DOI:** 10.1101/2021.11.15.468694

**Authors:** Mateusz Manicki, Halil Aydin, Luciano A. Abriata, Katherine A. Overmyer, Rachel M. Guerra, Joshua J. Coon, Matteo Dal Peraro, Adam Frost, David J. Pagliarini

**Author notes:** These authors contributed equally: Mateusz Manicki, Halil Aydin.

## Abstract

Coenzyme Q (CoQ, ubiquinone) is a redox-active lipid essential for core metabolic pathways and antioxidant defense. CoQ is synthesized upon the mitochondrial inner membrane by an ill-defined ‘complex Q’ metabolon. Here we present a structure and functional analyses of a substrate- and NADH-bound oligomeric complex comprised of two complex Q subunits: the hydroxylase COQ7, which performs the penultimate step in CoQ biosynthesis, and the prenyl lipid-binding protein COQ9. We reveal that COQ7 adopts a modified ferritin-like fold with an extended hydrophobic access channel whose substrate binding capacity is enhanced by COQ9. Using molecular dynamics simulations, we further show that two COQ7:COQ9 heterodimers form a curved tetramer that deforms the membrane, potentially opening a pathway for CoQ intermediates to translocate from within the bilayer to the proteins’ lipid-binding sites. Two such tetramers assemble into a soluble octamer, closed like a capsid, with lipids captured within. Together, these observations indicate that COQ7 and COQ9 cooperate to access hydrophobic precursors and coordinate subsequent synthesis steps toward producing mature CoQ.

## Main

Quinones are central to most molecular energy transducing systems in organisms across all kingdoms of life^1^. Many of the quinones essential for biology are attached to hydrophobic moieties that keep them sequestered within biological membranes. Within the bilayer, they participate in electron transfer between proteins involved in diverse processes, including respiratory and photosynthetic complexes^2, 3^. Despite their profound roles in life, many aspects of quinone biosynthesis and trafficking remained undefined^4, 5^.

Coenzyme Q (CoQ, ubiquinone) is a quinone found within every eukaryotic membrane^6^ and that is best known for its key role in the mitochondrial electron transport chain^7^. CoQ comprises a redox-active head group that can carry electrons and an extremely hydrophobic isoprenoid lipid tail that drives its partitioning into the hydrophobic portion of the bilayer. In eukaryotes, the head group is derived from aromatic amino acids and imported into mitochondria where it becomes attached to the isoprenoid tail. The head group is then chemically modified by several enzymes, each of which must access intermediates that are likely buried within the membrane^4^. Interestingly, individual intermediates of the pathway are difficult to detect, suggesting that substrate channeling might occur between the biosynthetic enzymes (COQ proteins). Supporting this, recent work has begun to define a dynamic complex of COQ proteins (complex Q, or CoQ-synthome) that may act in a metabolon-like fashion^8–10^.

Due in part to its dynamic nature—and the fact that structural information on multiple individual COQ proteins has not been reported—complex Q has evaded detailed structural and biochemical characterization. Protein interaction maps in Saccharomyces cerevisiae and human cells, however, suggest the complex is comprised of at least seven proteins (COQ3, COQ4, COQ5, COQ6, COQ7, COQ8, COQ9)^11–13^. The complex architecture and subunit stoichiometry remain unknown, as does whether discrete sub-complexes exist. However, observations in yeast^14^, mice^15^, and humans^16^ have shown that a direct physical and functional relationship exists between COQ7, an iron-dependent hydroxylase, and COQ9, a lipid-binding auxiliary protein, and that mutations in either protein lead to the accumulation of the penultimate CoQ intermediate, demethoxy-coenzyme Q (DMQ). Mutations of these proteins result in neurological defects, decreased activity of the mitochondrial electron transport chain, reduced levels of ATP synthesis, and increased mitochondrial oxidative stress^16–23^.

Details of COQ7 and COQ9 cooperation are not well understood. Recently, we demonstrated that COQ9 is a membrane-binding protein capable of selectively engaging aromatic isoprene lipids^24^. This observation suggested that COQ9 could assist COQ7 in membrane association and/or in accessing a CoQ intermediate. However, it is still unclear how COQ proteins surmount hydrophobic barriers to access CoQ intermediates, which reside entirely within the bilayer. To explore these ideas, we co-expressed and co-purified an octameric COQ7:COQ9 complex and solved its atomic structure using cryoEM. The structures show that COQ7 binds COQ9 via an extended ferritin-like fold, binds a CoQ precursor, and binds its NADH cofactor along with additional lipids. Within this complex, COQ9—which, on its own, binds CoQ precursors robustly—is in the apo form, suggesting that it may have delivered this ligand to COQ7. The octameric complex consists of two COQ7:COQ9 tetramers held together through their hydrophobic faces by various small hydrophobic molecules. While the significance of the octameric state of the complex remains unclear, biochemistry and simulations suggest that the tetrameric halves of the octamer may bind and bend target membranes, to enable these proteins to access lipids within the bilayer. Together, our data provide an initial insight into how a metabolon-like CoQ complex could assist the biosynthesis of a hydrophobic molecule at the membrane.

## Results

### CryoEM structure of human COQ7

COQ7 is a membrane-bound di-iron protein and an enigmatic member of the ferritin family that has been recalcitrant to experimental structural characterization^25–27^. Here, using an E. coli expression system, we co-expressed human COQ7 (residues 39-217 with N-terminal GB1 tag^28^) together with its native partner, human COQ9 (residues 79-318 with an N-terminal His tag), and performed metal affinity and gel filtration chromatography. Our results revealed that COQ7 and COQ9 co-purify as a 1:1 complex with a molecular weight of ∼240 kDa, suggesting a likely octamer (Fig. 1a and Extended Data Fig. 1a-d). Further biophysical characterization by negative-stain electron microscopy confirmed the homogeneity of the sample and revealed monodispersed particles of the intact COQ7:COQ9 complexes with symmetric features (Extended Data Fig.1e,f).

**Figure 1.**
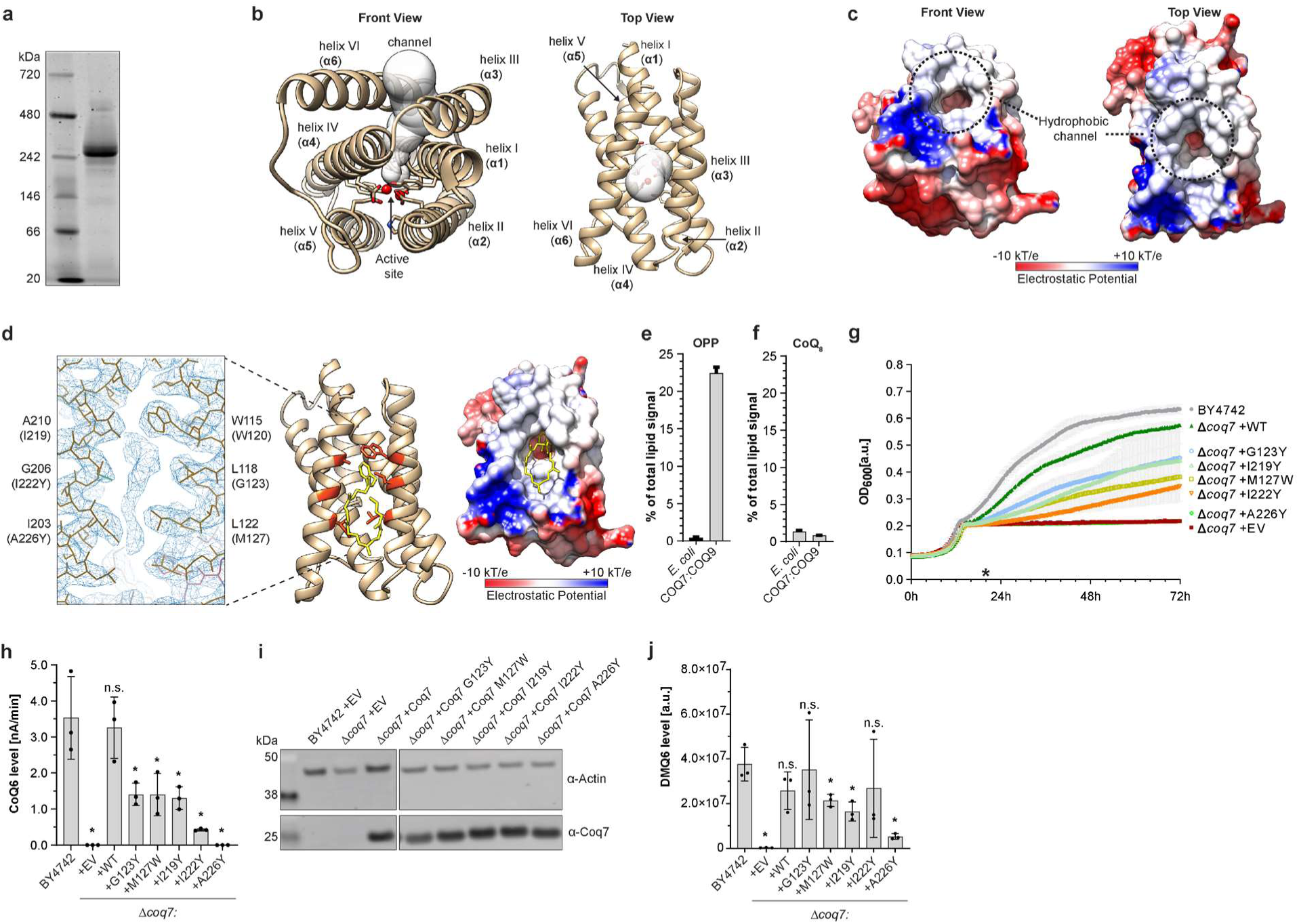
Structure – function characterization of COQ7. **a**, Native PAGE analysis of the homogeneity and size of purified GB1-^Nd38^COQ7:His6- ^Nd79^COQ9 complex. **b**, Structure of COQ7 with visible channel leading to an active site. Iron atoms were not resolved by cryoEM and were added based on the structure of bactoferritin (PDB:4AM2). **c**, Electrostatic surface potential of COQ7 with visible entry to the hydrophobic channel. **d**, Detailed view of the hydrophobic channel with visible additional ligand density inside. Important residues are labeled. Corresponding residues in yeast are in parentheses. **e,f,** LC-MS lipidomic analysis of relative levels of Octaprenylphenol (OPP) and Coenzyme Q8 (CoQ8) in E. coli cells and the GB1-^Nd38^COQ7:His6-^Nd79^COQ9 complex purified from E. coli. (mean ± s.d., n = 2) **g**, Respiratory growth of wild type BY4742 yeast and Coq7 mutants expressed in BY4742 Δcoq7 genetic background. Cells start with fermentation and then switch to respiratory growth (mean ± s.d., n = 3). The star (*) corresponds to time point used for measurements in panel h,i,j. **h**, HPLC-ECD analysis of Coenzyme Q6 level in wild type BY4742 yeast and Coq7 mutants expressed in BY4742 Δcoq7 genetic background. Signal normalized to internal CoQ8 control (mean ± s.d., n = 3, one-sided Student’s t-test, *p<0.05, n.s. not significant). i, Expression level of the Coq7 yeast mutants (p413, TEF promoter) measured with Western Blot. Native Coq7 in BY4742 strain is below the limit of detection. Samples were prepared in triplicates. Representative membrane shown. **j)** LC-MS lipidomic analysis of DMQ6 substrate level in wild type BY4742 yeast and Coq7 mutants expressed in BY4742 Δcoq7 genetic background. Signal normalized to internal CoQ8 control (mean ± s.d., n = 3, one-sided Student’s t-test, *p<0.05, n.s. not significant).

The complex was vitrified, imaged, and reconstructed using 3D single-particle cryoEM analysis. An initial density map reached ∼3.5Å but was incompletely occupied by the known cofactor NADH (Extended Data Fig. 2-3, Extended Data Table 1). We therefore incubated the COQ7:COQ9 hetero-octamers with excess NADH prior to vitrification. This second dataset yielded an improved map at an overall nominal resolution of ∼2.4 Å, with D2 symmetry, and robust density for NADH (Extended Data Fig. 4-5, Extended Data Table 1). Most of the side chains were well-resolved and allowed de novo atomic model building of COQ7 and COQ9, with the exception of the flexible N-terminal GB1 tag on COQ7 and C-terminal 10^th^ α-helix of COQ9 (Extended Data Fig. 6, Extended Data Table 1).

COQ7 has a rod-shaped structure (Fig. 1b) with approximate dimensions of 45 Å X 30 Å X 27 Å and shares structural similarity with bacterioferritin (∼1.7 Å r.m.s.d. over 173 Cα atoms of COQ7) (Extended Data. Fig. 7a). However, major structural changes are apparent and suggest that the fold was repurposed from the classic iron-storing ferritin to a membrane-binding hydroxylase (Extended Data Fig. 7b). The overall structure is comprised of six α-helices (α1- α6) (Fig. 1b). Helices α1, α2, α4, and α5 exhibit a compact four-helix bundle containing the di-iron center, which is able to bind two metal ions (Fe1 and Fe2). The three-dimensional map of COQ7 presents the atomic coordinates of the structurally and functionally important iron coordination motif (E-(X6)-Y-(X22)-E-(X2)-H-(X48)-E-(X6)-Y-(X28)-E-(X2)-H), which is conserved across di- iron carboxylate proteins, supporting the enzymatic function of the protein as a hydroxylase (Extended Data Fig. 7c). While the protein chains were well defined by electron density, we did not observe electron density for the active site ions and could only estimate their position. By analogy with bacterioferritin, the first metal ion binding site, Fe1, would be coordinated by E60, H93, Y149, and E178, whereas the Y67, E90, E142, and H181 residues would coordinate the Fe2 site.

In addition to the four-helix bundle, COQ7 structure contains a short helix (α3) and a C-terminal helix (α6) that is connected to the rest of the molecule via a long loop. These features represent a novel organization for ferritin-like di-iron carboxylate enzymes (Extended Data Fig. 7d). The helices α3 and α6 contribute to the formation of a large hydrophobic channel along with α1 and α4 (Fig. 1b and Extended Data Fig. 7e), and a flat hydrophobic surface on the membrane-proximal side of the protein (Fig. 1c). The residues L111, L114, L118, L122, I202, V209, L213, and L217 define the outer entrance to the channel, whereas the inner surface is lined by A63, I66, W115, V141, I145, I203, C207, A210, and I211. This hydrophobic surface may interact with the surface of the mitochondrial inner membrane, and the channel is sufficiently large to admit the entry of quinone intermediates and access to the di-iron center. A similar structural feature can be found in alternative oxidases (AOX) that also belong to the ferritin family and that use CoQ as a substrate for different roles in the cell^29^ (Extended Data. Fig. 7f-h).

We also observed a clearly defined isoprenoid quinone-like density extending from the inner surface of the hydrophobic channel and making several contacts with the hydrophobic surface (Fig. 1d and Extended Data Fig. 6d). The quinone head group fits comfortably into the hydrophobic pocket and the isoprene tail adopts a U-shaped conformation that is stabilized by conserved hydrophobic residues (L118, A121, L122, L129, L199, and I202) located on the protein surface. To determine the identity of this isoprenoid, we performed liquid chromatography with tandem mass spectrometry (LC-MS/MS) of the purified COQ7:COQ9 complex and of bacterial cells used for its production. The protein sample showed 50x enrichment (∼20% of total lipid signal) in octaprenylphenol (OPP), a bacterial CoQ biosynthetic intermediate^30^ (Fig. 1e). In agreement with the logic of biosynthetic pathways, no enrichment of CoQ8 (the E. coli version of CoQ) was detected showing specificity of the proteins towards the CoQ intermediate and exclusion of the mature CoQ product (Fig. 1f). Together, these results identify OPP as the bacterial ligand occupying the hydrophobic channel leading to the active site of COQ7.

To functionally validate these observations, we turned to yeast that can survive without CoQ in fermentation but require it for respiration. Structural features and evolutionary conservation of COQ7 (Extended Data Fig. 8a) guided us in the design of mutations that should block the hydrophobic channel. As expected, mutations G123Y, M127W, I219Y, I222Y, or A226Y (corresponding to human L118, L122, I203, G206, or A210 residue) that introduce a large hydrophobic residue into the channel, reduced CoQ6 levels in yeast and their ability to respire (Fig. 1g, h) without affecting the proteins’ stability as judged by their expression levels (Fig. 1i). To further confirm specific disruption of Coq7’s function in the yeast mutants we used mass-spectrometry to measure levels of DMQ6, the bona fide substrate of Coq7. In all cases, we detected the molecule showing that upstream steps of CoQ6 biosynthesis are at least partially functional and that low Coq7 activity may constitute a pathway bottleneck in the mutant strains (Fig. 1j). Together, these data strongly support in vivo existence of the hydrophobic channel, as evidenced by the COQ7 structural model. Of note, the structure helps to understand the deleterious effects of COQ7 mutations found in human patients, including a recent case of K200fs56 mutation^17^ that localizes to COQ7’s 6^th^ helix revealed for the first time in this manuscript (Extended Data Fig. 8b).

### CryoEM reveals details of NADH binding by COQ7

To achieve their enzymatic activity, di-iron proteins require a reduction of the iron atoms in their active site from Fe^3+^ to Fe^2+^ state^31^. It has been shown in vitro that quinone substrates can mediate the transfer of electrons from NADH to COQ7’s active site^32^. Comparison of our unsoaked and NADH-soaked reconstructions showed how NADH binds to COQ7 (Fig. 2a) via a set of evolutionarily conserved surface-exposed residues (Extended Data Fig. 8a) without inducing global conformational changes. Although it is unknown how the charge is transferred from the NADH to the protein, it has been suggested that it might follow an uncommon, sequential electron-proton-electron donation mechanism^26^, and involve an amino acid triad E60/H148/Y149^32^. In our structure the cofactor localizes next to a water-filled channel formed by the triad that might facilitate proton transfer. It is also close enough to the active site pocket (∼15 Å) to permit direct tunneling of electrons to the expected quinone acceptor^33–36^ (Fig. 2b,c and Extended Data Fig. 9a-d,i). This biologically useful positioning of NADH is provided by the R51, R208, Y212, and R216 residues of COQ7 that contact different parts of the NADH molecule (Fig. 2d). Interestingly, the interaction between R208 and NADH is bridged by the headgroup of a clearly visible, but not identified, phospholipid molecule (Fig. 2d). This suggests a functional role of membrane lipids in the stabilization of NADH contacts.

**Figure 2.**
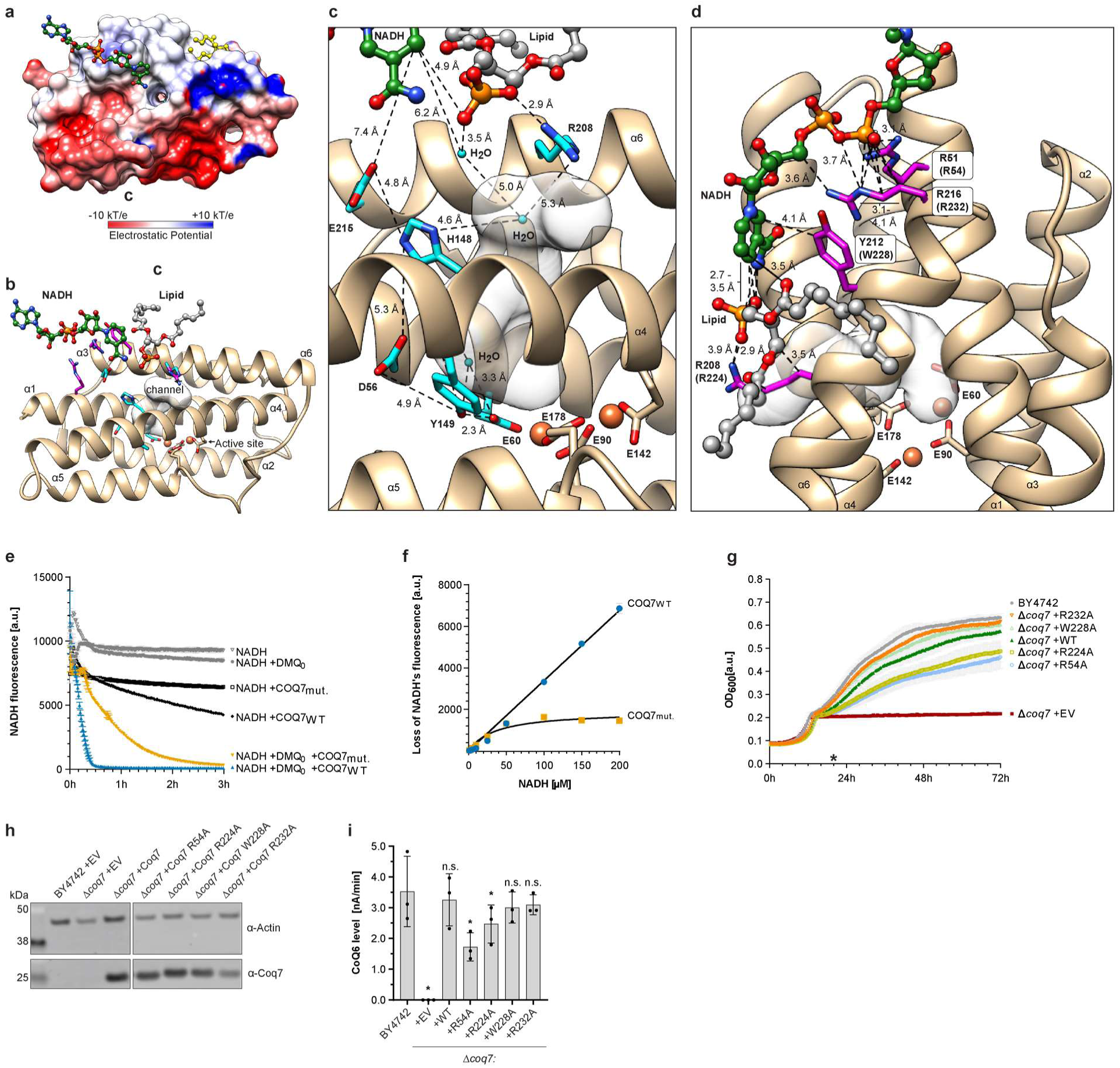
Details of NADH binding to COQ7. **a**, Electrostatic surface potential of COQ7 with bound NADH molecule (green) and OPP (yellow). **b**, Ribbon view of COQ7’s NADH binding site and channel leading to the active site. **c**, Structural elements that might participate in charge transfer from NADH to COQ7. **d**, Details of NADH binding by conserved arginines and tyrosine. **e**, NADH oxidation in vitro assay showing activities of WT and RRYR/AAAA quad COQ7 mutant (mean ± s.d., n = 3). **f**, Effect of the RRYR/AAAA mutation as a function of NADH concentration. The graph shows 30 min. time point extracted from data presented in Extended Data. Fig. 9g,h (mean ± s.d., n = 3). **g**, Respiratory growth of wild type BY4742 yeast and Coq7 mutants expressed in BY4742 Δcoq7 genetic background. Cells start with fermentation and then switch to respiratory growth (mean ± s.d., n = 3). The star (*) corresponds to time point used for measurements in panel h,i, **h**, Expression level of the Coq7 yeast mutants (TEF promoter) measured with Western Blot. Native Coq7 in BY4742 strain is below the limit of detection. Samples were prepared in triplicates. Representative membrane shown. i, HPLC-ECD analysis of Coenzyme Q6 (CoQ6) level in wild type BY4742 yeast and Coq7 mutants expressed in BY4742 Δcoq7 genetic background. Signal normalized to internal CoQ8 control (mean ± s.d., n = 3, one-sided Student’s t-test, *p<0.05, n.s. not significant).

To test the functional importance of the identified residues, we purified COQ7 alongside a quadruple COQ7^R51A/R208A/Y212A/R216A^ mutant (Extended Data Fig. 9e) and tested their activity in vitro by measuring the loss of NADH fluorescence upon its oxidation to NAD+. In agreement with its proposed mechanism of action, wild type COQ7 slowly oxidized NADH and transferred electrons to a quinone acceptor, changing its redox state as measured by an HPLC equipped with a redox-sensitive electrochemical detector (HPLC-ECD) (Extended Data Fig. 9f). However, the mutant markedly lost activity (Fig. 2e,f and Extended Data Fig. 9g,h). Although arginines have been reported to play a part in NADH binding (e.g., Rex^37^, p-Hydroxybenzoate Hydroxylase^38^, Malic Enzyme^39^) they are not the most common residues found in NADH-binding motifs^40, 41^. Therefore, we again sought to validate the biological importance of our observations in vivo using the yeast respiration system. High overexpression of Coq7^R54A^ or Coq7^R224A^ point mutants (corresponding to residues R51 and R208 in human COQ7, respectively) was not able to fully support CoQ6 production or respiratory growth. Interestingly, mutants W2228A and R232A (Y212 and R216 in humans) showed only a minor reduction in CoQ6 level, and hence fully supported respiration (Fig. 2g-i). The partial loss of the in vitro activity of the quadruple human COQ7 mutant and the reduced levels of CoQ6 in yeast mutants indicate that the NADH’s positioning observed in cryoEM is biologically relevant. Furthermore, as no structure of COQ7 has yet been reported, our work provides a first glimpse into how this protein leverages its fold to perform the penultimate step of CoQ biosynthesis.

### CryoEM reveals lipid molecules bound to the COQ7:COQ9 heterodimer

Efficient interaction of COQ7 with COQ9 is known to be essential for CoQ production in vivo. We previously described the crystal structure and lipid-binding activity of human COQ9 and used molecular dynamics simulations to predict its interaction with a COQ7 homology model on a lipid membrane^24^. Here, using cryoEM, we experimentally determined the structure of the complex bound to lipids. The atomic structure of human COQ9 (α1 to α9) fits well as a rigid body to the electron density of the COQ7:COQ9 complex. Interestingly, the α7- α8 loop adopts only one of the two conformations found in the isoprene-bound crystal structure of COQ9 (PDB:6DEW). The overall architecture of the COQ7:COQ9 heterodimer is shown in Fig. 3a. We observed an interface that is stabilized by several hydrogen bonds, hydrophobic interactions, and an intermolecular salt bridge between COQ9 (D225, D226, T236, D237, F238, W240, and Y241) and COQ7 (N98, M101, V102, R105, R107, P108, T109, V110, and M112) residues, and that spans a buried solvent-accessible surface area of approximately 468 Å^2^ (Fig. 3b). Notably, this limited contact interface between COQ7 and COQ9 may not be sufficient for stable complex formation. In fact, a single point mutation in that region COQ9^W240K^ led to total disruption of the octameric 240 kDa COQ7:COQ9 complex and formation of a smaller ∼55 kDa complex comprised only of COQ9 (Fig. 3c and Extended Data Fig. 10a-c). This shows that W240 is crucial for this contact site between COQ7 and COQ9, which ultimately leads to formation of a higher mass complex stabilized by lipids, isoprenes and NADH.

**Figure 3.**
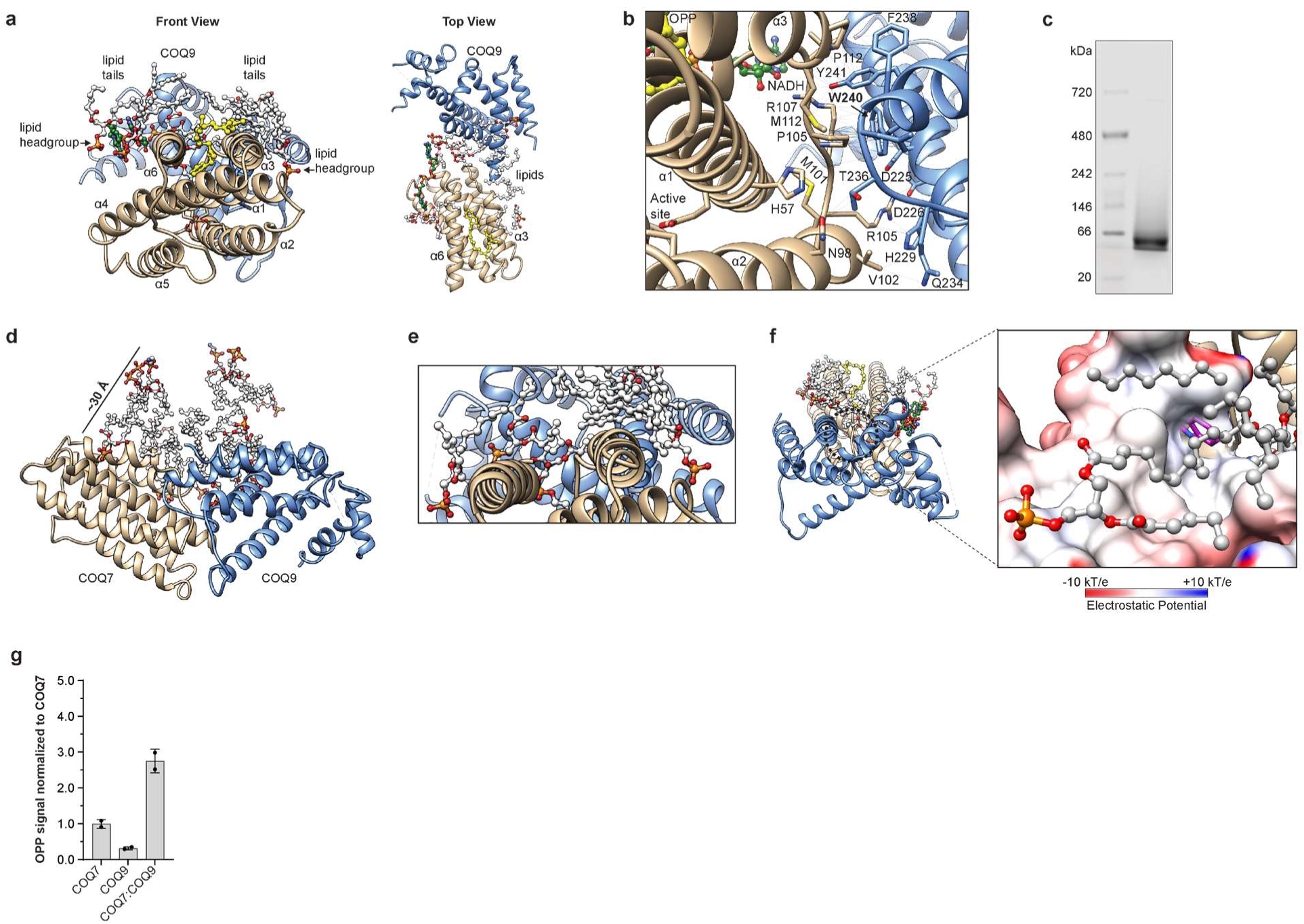
Structure of COQ7:COQ9 heterodimer in lipid environment. **a**, Snapshot of the COQ7:COQ9 heterodimer structure with visible lipids (light gray), NADH (green) and OPP (yellow). **b**, Details of the COQ7:COQ9 interaction interface. **c**, Native Page of the mutated GB1-^Nd38^COQ7:His6-^Nd79^COQ9^W240K^ complex. **d**, Zoom on lipid molecules forming a lipid pseudo-bilayer. **e**, Zoom on the tilted lipid headgroups (orange and red). **f**, Zoom on lipid-binding substrate pocket in COQ9 surrounded by surface lipids. No quinone density is detected around the substrate-binding COQ9’s W240 residue (magenta). **g**, LC-MS lipidomic analysis of OPP content in individually purified His6-GB1-^Nd38^COQ7, His6-^Nd79^COQ9 or the full GB1- ^Nd38^COQ7:His6-^Nd79^COQ9 complex (mean ± s.d., n = 2).

In particular, investigation of the 2.4 Å cryoEM density map revealed multiple lipid-like densities engaged with the surface of the proteins. They were not randomly oriented, but rather formed a 30 Å tall, bilayer-like and warped sheet with lipid tails pointing towards each other and the headgroups pointing in opposite directions (Fig. 3d). The lipids likely originate from E. coli plasma membrane based on our lipidomics results^42, 43^ (Extended Data Fig. 10d). These observations may have biological relevance because quinones are known to be trapped within the membranes. Due to their extreme hydrophobicity, these molecules cannot transverse through the headgroup layer^44–49^ and thus likely have to be extracted by proteins.

We previously determined the crystal structure of COQ9 with a hydrophobic isoprenoid substrate and identified a lipid-binding pocket inside the protein. In the cryoEM structure, this pocket is located exactly between the displaced lipid headgroups, consistent with a model in which COQ9’s binding reorganizes the membrane to enable access to its interior. Interestingly, we did not observe a well-defined electron density for lipids resembling CoQ intermediates within the lipid-binding cavity of COQ9 (Fig. 3f) which could indicate that the quinone molecule was transferred to COQ7. If that is the case, then COQ7 should be enriched in OPP when in complex with COQ9. To verify this, we measured quinone content in COQ7 and COQ9 purified alone and as a complex. As expected, the lipidomic analysis confirmed a higher OPP level in the COQ7:COQ9 complex than in COQ7 or COQ9 alone (Fig. 3g and Extended Data Fig. 10e). Knowing there is no observable OPP bound to COQ9 in the COQ7:COQ9 complex, the higher OPP signal most likely represents an increased level of OPP bound to COQ7 due to COQ9’s actions. Together, our structural model reinforces and advances our previous in vitro analyses, and provides a detailed blueprint for defining COQ7-COQ9-lipid interactions critical for complex functionality.

### COQ7 and COQ9 form a heterotetramer

It has been reported that in solubilized human mitochondria COQ7 is predominantly detected as a complex of 100-150 kDa mass^50^. A separate study found COQ9 to migrate in 50-150 kDa range^51^. This suggests that a subcomplex of our ∼240 kDa human COQ7:COQ9 octamer may represent its dominant in vivo state. In the cryoEM structure, two heterodimers of COQ7:COQ9 come together on the same side of the lipid pseudo-bilayer and form a curved tetramer with a hydrophobic surface suitable for membrane interaction. Viewed from the membrane surface, one heterotetrameric assembly is approximately 100 Å in diameter and 40 Å axially with a solvent-exposed central pore that accommodates the NADH molecule (Fig. 4a). It is formed by two COQ7:COQ9 heterodimers that make additional contacts with an interface that buries around 261 Å^2^ of surface area. The N-terminal segment of the α7 helix region of COQ9 and α6 helix of COQ7 from opposing heterodimers associate around a 2-fold axis normal to the membrane plane, with most intimate contacts occurring between COQ9 residues L209, P210, and H211 and COQ7 residues S201, Q204, A205, and R208 (Fig. 4b).

**Figure 4.**
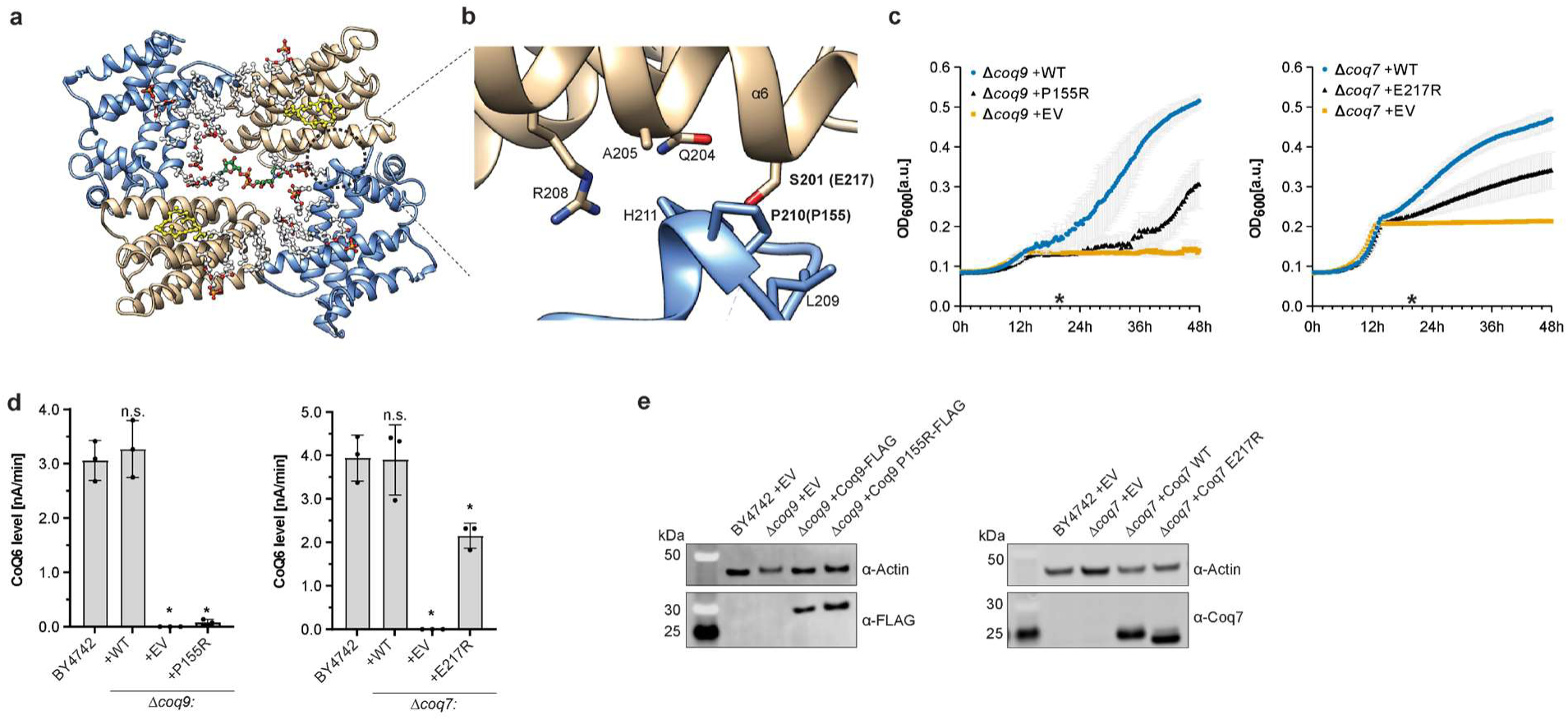
Structure of the COQ7:COQ9 heterotetramer. **a**, Membrane view of an individual COQ7:COQ9 tetramer. **b**, Details of COQ7:COQ9 interaction mediating tetramer’s formation. COQ9 P210 and COQ7 S201 residues are labeled in bold and their corresponding residues in yeast are provided in parentheses. **c**, Respiratory growth of Coq7 or Coq7 mutants expressed in BY4742 Δcoq7/Δcoq9 genetic background. Cells start with fermentation and then switch to respiratory growth (mean ± s.d., n = 3). The star (*) corresponds to time point used for measurements in panel d,e. **d**, HPLC-ECD analysis of Coenzyme Q6 level in wild type BY4742 yeast and Coq7 or Coq9 mutant expressed in BY4742 Δcoq7 or Δcoq9 genetic background. Signal normalized to internal CoQ8 control (mean ± s.d., n = 3, one-sided Student’s t-test, *p<0.05, n.s. not significant) e, Expression level of the Coq7 yeast mutants (TEF promoter) measured with Western Blot. Native Coq7 in BY4742 strain is below the limit of detection. Samples were prepared in triplicates. Representative membrane shown.

To test the importance of this region, we generated corresponding yeast mutants and measured their CoQ6 levels and ability to respire. We observed that mutations of P210 and S201—represented by Coq9^P155R^ and Coq7^E217R^ in yeast— led to impaired respiration and diminished CoQ6 levels, despite high expression levels of the mutated proteins (Fig. 4c-e). Over time, yeast did display moderate respiratory growth showing that the proteins remain at least partially functional. Given the magnitude of the Coq9^P155R^ phenotype, we queried whether disruption of that residue disturbs the human COQ7:COQ9 complex. The COQ9^P210K^ mutation did not prevent the formation of the 240 kDa complex expressed in bacteria at 20°C (Extended Data Fig. 10f) suggesting that the interaction site is not fully disrupted or that protein-protein interactions are not the sole contributors to its stability. Due to the large hydrophobic surfaces present in COQ7 and COQ9, we hypothesize that membrane association may play an important role in that stabilization.

### Molecular dynamics simulations of the tetramer engaging and shaping a membrane

To explore whether and how the tetrameric form binds the lipid bilayer to potentially facilitate transformation of the embedded quinone substrate, we performed atomistic molecular dynamics (MD) simulations on systems set up with the COQ7:COQ9 heterotetramer manually placed with its hydrophobic surface facing but not binding a model mitochondrial membrane. The systems were parametrized as described in methods. Of note, since no parameterized models of the quinone detected in the cryoEM structure (OPP) or the true COQ7:COQ9 substrate (DMQ) are available, we used CoQ10 by Galassi et al.^52^ as an approximation of the small molecules involved in COQ7 and COQ9 activities. CoQ10 was either added to the membrane at ∼4% concentration in simulations designed to test how the substrate-free protein binds to membranes, or placed individually inside COQ7’s active site in simulations aimed at testing its stability when bound to COQ7’s active site.

In all simulations, the tetrameric arrangement inferred from the cryoEM structure interacted rapidly (<50 ns) with the membrane. This interaction remained stable for the whole simulation (Fig. 5a). Upon this interaction, COQ7 progressively gained direct access to the hydrophobic portion of the membrane (plotted as contacts between COQ7 and the carbon tails of the lipids over time in Figure 5a). As COQ7’s access to the membrane proceeded, lipids tilted away from a normal bilayer architecture until they resembled the angular orientations of the lipids resolved in the cryoEM density map. In addition, the lipids concentrated their headgroups on the periphery of the protein-membrane contact region to form a rim around the membrane-facing surface of the COQ7 proteins (Fig. 5b). This arrangement exposed the carbon- tail region of membrane lipids directly to the COQ7 proteins (Fig. 5b), while leaving COQ9 to interact peripherally with the headgroup phosphates. As a consequence, a strong local deformation of the membrane of around 10Å was observed (Fig. 5c and Extended Data Fig. 11a,b).

**Figure 5.**
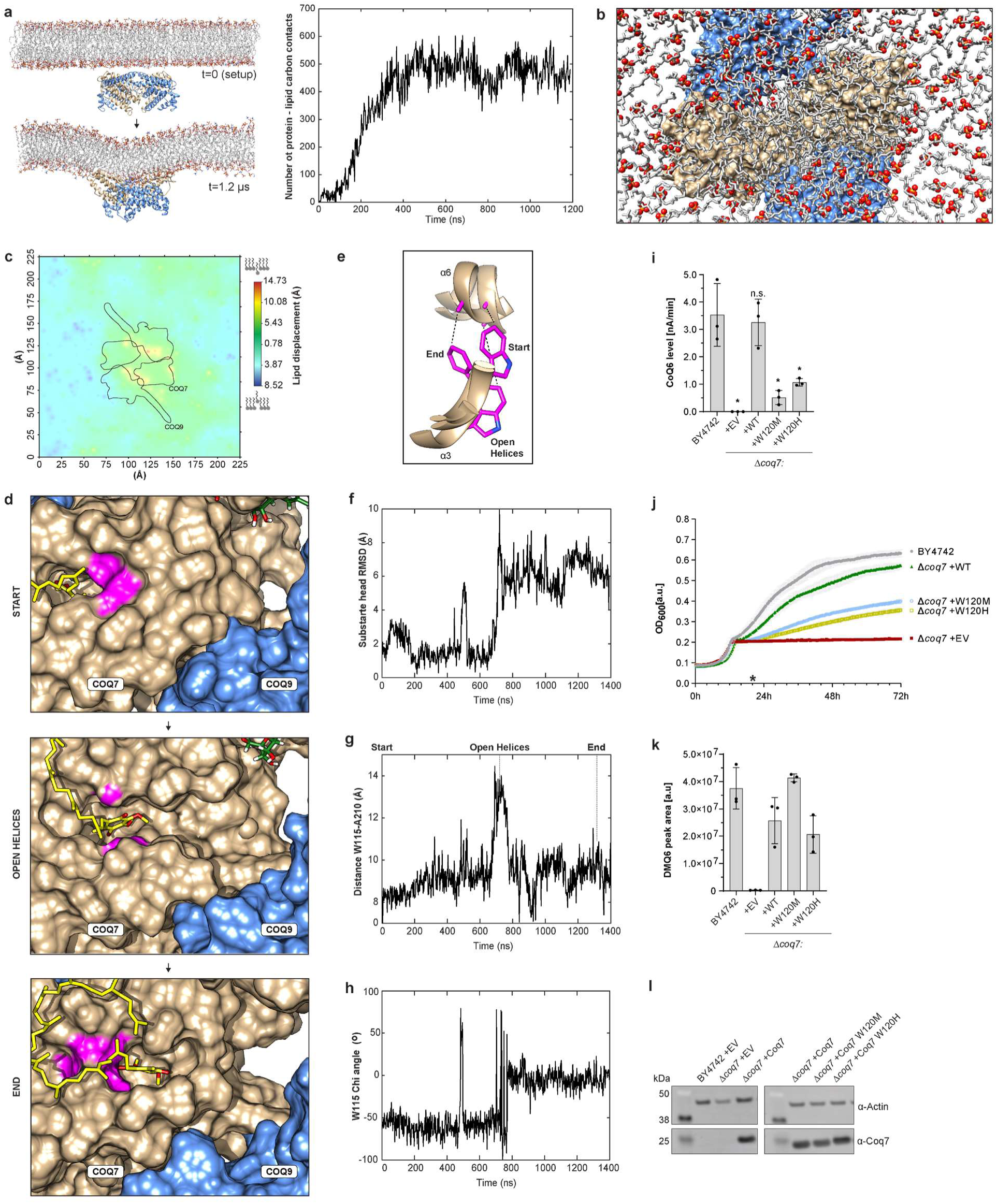
Molecular dynamics simulations of the COQ7:COQ9 tetramer at the membrane. **a**, Membrane binding and deformation observed in the atomistic MD simulations. The graph describes the number of contacts between COQ7 atoms and carbon atoms of all lipid tails. **b**, Shot of the protein-membrane complex observed after a few hundreds of ns of simulation when the tetramer has anchored to the membrane, exemplified here with the end point of one trajectory. Phosphate groups representing lipid headgroups are displayed as red spheres. Other atoms are displayed as sticks. COQ7 (tan) and COQ9 (blue) are visible as surfaces. Notice how COQ7 gains direct access to the hydrophobic portion of the membrane. **c**, Map of the deformation of the bottom membrane leaflet induced by the proteins binding to it. **d**, Snapshots from molecular dynamics simulation at 450 K showing release of CoQ10 from COQ7 (tan) towards COQ9 (blue) by gating through W115 and A210 (magenta). **e**, RMSD of the CoQ10 molecule as a proxy for its displacement out of the active site. **f**, Overlaid positions of the gating W115 and A210 residues from COQ7 conformations visible in panel d. **g**, Time plot showing the distance between the Cα atoms of W115 and A210. **h**, Time plot showing the sidechain torsion angle of W115. **i**) LC-MS lipidomic analysis of DMQ6 level in wild type BY4742 yeast and Coq7 mutants expressed in BY4742 Δcoq7 genetic background. Signal normalized to internal CoQ8 control (mean ± s.d., n = 3, one-sided Student’s t-test, *p<0.05, n.s. not significant). **j**, HPLC-ECD analysis of Coenzyme Q6 (CoQ6) level in wild type BY4742 yeast and Coq7 mutants expressed in BY4742 Δcoq7 genetic background. Signal normalized to internal CoQ8 control (mean ± s.d., n = 3, one-sided Student’s t-test, *p<0.05, n.s. not significant). **k**, Respiratory growth of wild type BY4742 yeast and Coq7 mutants expressed in BY4742 Δcoq7 genetic background. Cells start with fermentation and then switch to respiratory growth (mean ± s.d., n = 3). The star (*) corresponds to time point used for measurements in panels i,j,l. l, Expression level of the Coq7 yeast mutants (TEF promoter) measured with Western Blot. Native Coq7 in BY4742 strain is below the limit of detection. Samples were prepared in triplicates. Representative membrane shown.

Given the direct contact between the hydrophobic surfaces of COQ7 and the internal, hydrophobic portion of the membrane, we speculate that the local membrane reshaping induced by the tetramer might help to facilitate the exchange of substrates and products directly from the membrane into and out of the tetrameric complex. Indeed, in the simulations with model membranes doped with 4% CoQ10 we observed transient association of the small molecule with the hydrophobic surface of COQ7 (Extended Data Fig. 11c). However, the aromatic headgroup of CoQ10, which has similar size to COQ7’s bona fide substrate DMQ, seemed too bulky to access the protein’s active site (Extended Data Fig. 11d). Therefore, its entry may require a conformational change of COQ7 which is unlikely to spontaneously take place in microsecond-long atomistic simulations.

To explore this idea, we devised a reversed computational experiment in which we modeled a molecule of CoQ10 inside COQ7’s active site based on the position of OPP molecule observed in the cryoEM structure, and then monitored its release from the active site through simulations. Replica-exchange simulations were impossible to tune, as they resulted in membrane disruption; we therefore tested regular simulations at various stable temperatures. At 450 K we observed events in which the docked CoQ10 dissociated from the active site without compromising the protein or membrane stabilities within the simulated timescale (∼1-1.5 microsecond).

Out of 10 replicas at 450 K, we observed CoQ10 leaving the active site in two directions: 8 times directly towards and into the membrane bulk, and twice towards COQ9 before moving to the membrane bulk (Fig. 5d and Extended Data Fig. 11e). In every case, release of the ligand involved a displacement of the two COQ7 helices (α3, α6) that form the hydrophobic channel leading to the active site, temporarily widening its opening towards the membrane.

Although it is hard to tell how much of this is due to the high temperatures used to promote unbinding and how much reflects realistic motions, we noticed that the residue W115 flips its sidechain a few times (Fig. 5e) in the pathway where the CoQ10 molecule moves towards COQ9 (Fig. 5f). This is realized through rotation of the Chi1angle (Fig. 5g) concerted with separation of the two helices represented by the increase of the distance between W115’s and A210’s Cα atoms (Fig. 5h). Once the small molecule has left the active site, W115’s sidechain flips to a second stable conformation where the indole ring fills the space that the headgroup of CoQ10 was occupying (Fig. 5f,h). We thus postulated that residue W115 could act as a gate that communicates a potential pathway of substrate entry from an initial binding site around COQ9 with the active site at COQ7, and possibly also modulate separation of COQ7’s helices to allow release of the product towards the membrane.

To address the relevance of this tryptophan experimentally, we mutated the corresponding W120 residue in yeast. Substitution with much smaller but also hydrophobic methionine decreased CoQ6 levels (Fig. 5i), compromised respiration (Fig. 5j) but mostly preserved levels of the Coq7’s bona fide substrate DMQ6 (Fig. 5k). High expression level of the mutant (Fig. 5l) indicates that the Trp residue is indeed important for the function of COQ7 and not its stability. Accordingly, the substitution of W120 in yeast with histidine, which also contains an aromatic ring but is more polar than the tryptophan, also distorted the production of CoQ6 (Fig. 5i-l).

Why the activity of COQ7 depends on a specific aromatic ring remains unclear. However, the aromatic nature of its quinone substrate suggests that π interactions might play a role in substrate selection and loading, possibly facilitated by COQ9 and its essential quinone- binding W240 residue. At the same time, COQ7’s structural similarity with alternative oxidase, a protein whose function is to oxidize mature CoQ, suggests that COQ7 might require an additional layer of control to promote hydroxylation of the DMQ intermediate and avoid oxidation of the final CoQ product.

Our MD system offers novel insights and a framework for future studies of how the COQ7:CO9 binds the membrane, modify its structure, and delivers a quinone intermediate to COQ7.

### The overall COQ7:COQ9 complex forms a hetero-octameric cage

Physiologically, COQ proteins are associated with the matrix side of the inner mitochondrial membrane^53, 54^ where they can interact with each other to produce CoQ. The single-leaflet- binding heterotetrameric COQ7:COQ9 fits well into this model. However, the cryoEM revealed that the purified COQ7:COQ9 complex forms a soluble ∼240 kDa capsid-like octamer composed of two heterotetramers that associate via contacts between the helices α7, α8, and α9 of COQ9 and α3 and α6 of COQ7 (Fig. 6a). The functional significance of this assembly state remains unclear.

**Figure 6.**
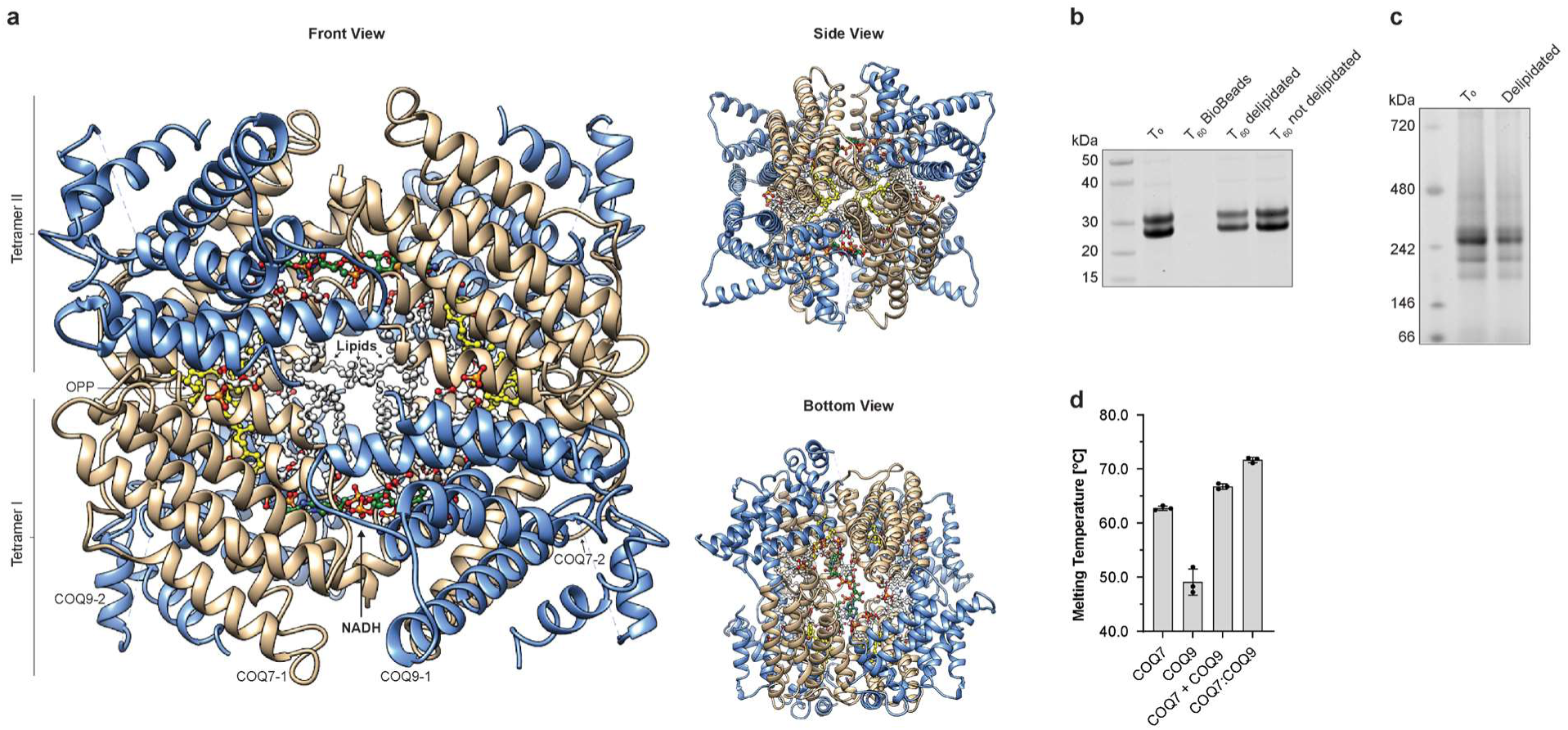
Two COQ7:COQ9 tetramers form an octamer. **a** Structure of the octamer with visible bound NADH (green) and OPP (yellow) **b**, SDS-PAGE analysis of protein stability upon delipidation of the GB1-^Nd38^COQ7:His6-^Nd79^COQ9 with BioBeads; T_0_ - soluble fraction before delipidation, T_60_ BioBeads - protein bound to beads after 60 min. incubation, T_60_ delipidated - soluble fraction after 60 min. delipidation, T_60_ not delipidated - soluble fraction after 60 min. incubation without BioBeads. **c**, Native PAGE analysis of the GB1-^Nd38^COQ7:His6-^Nd79^COQ9 complex after 15 min. delipidation. **d**, DSF analysis of melting temperatures of individually purified His6-^Nd38^COQ7, His6-^Nd79^COQ9, their mix and the GB1-^Nd38^COQ7:His6-^Nd79^COQ9 complex.

Globally, the octamer resembles a cube with a proteinaceous outer shell, a middle layer of protein-interacting lipids, and an inner core consisting of clearly visible densities. We identified twelve phospholipid densities occupying octamer interfaces. Our lipidomics revealed that highly abundant phospholipid species in E. coli copurified with recombinantly expressed COQ7 and COQ9 (Extended Data Fig. 10d). When considered with the structure (Fig. 3d), and the in silico simulations of membrane binding (Fig. 5a), these results suggest that lipids shield the hydrophobic surfaces of the proteins and therefore assist solubility and stability.

To test this notion, we delipidated the sample using lipid-capturing BioBeads and monitored soluble complex concentrations using SDS-PAGE gels. We observed 33% loss of the proteins upon 1h incubation (Fig. 6b) without the appearance of any new oligomeric structures, as judged by Native PAGE analysis (Fig. 6c). This indicates that specific protein-lipid interactions are important for the stability of the COQ7:COQ9 complexes isolated from E. coli. In addition, we asked if protein-protein interactions also have a stabilizing effect. Therefore, we performed thermal denaturation assays to measure the stability of the lipid-rich but individually purified COQ7, COQ9, COQ7 and COQ9 combined in vitro, and the COQ7:COQ9 complex.

The Tm of the wild-type COQ9 and COQ7 were 49.1 °C and 62.7 °C, respectively. Mixing of the proteins increased the melting temperature to 66.8°C which shows that COQ proteins are strongly stabilized by direct interactions in vitro. The melting temperature of the purified COQ7:COQ9 complex was higher (71.6°C) possibly representing a more optimal organization of lipids and proteins in the complex (Fig. 6d).

The octameric structure of the COQ7:COQ9 complex with the membrane trapped inside sparks intriguing ideas about its potential roles, for example as a lipid transporter or an enzymatic nano-chamber. Such functional states are rather unexpected, since the preponderance of evidence places COQ proteins only on the matrix face of the IMM^53–56^. However, in human and nematodes COQ7 has been reported to exist also in the nucleus^57^. Although these findings are debated^58^, they highlight that COQ7 may have additional functions, and thus the COQ7:COQ9 octamer warrants further experimental investigation. Collectively, we conclude that our data argue in favor of the COQ7:COQ9 tetramer being a physiologically-active form of the complex that can engage lipid bilayers and whose formation depends on protein-protein and protein-lipid interactions. The significance of the octameric state will be addressed in future studies.

## DISCUSSION

Quinones are highly reactive molecules^59^ that can be produced by cells in the form of lipids and employed as essential redox cofactors for membrane-bound enzymes^3, 60^. However, the biosynthesis of these hydrophobic quinones presents known biochemical challenges at the aqueous/membrane interface^61^. Enzymes involved in this biosynthesis require simultaneous access to water-soluble cofactors and lipid-soluble substrates and need to avoid the release of immature precursors into the membrane that may lead to toxic interactions with the quinone-dependent enzymes^62^. In the CoQ pathway, prenylation occurs at a very early step^4^, thereby creating a scenario whereby all subsequent intermediates are poised to partition into the membrane. How the enzymes circumvent this problem is largely unknown.

In recent years, evidence for a complex of CoQ-related biosynthetic proteins has emerged^8, 9, 12, 14, 50, 63–68^, leading to the speculation that CoQ is produced by a type of ‘metabolon’ that might surmount these challenges. Metabolons are dynamic protein clusters that create dedicated compartments to facilitate substrate channeling and protection of toxic or labile intermediates^69^. Indeed, in E. coli, multiple proteins in the CoQ pathway form a soluble complex in the cytosol that completes a series of enzymatic reactions before returning to membrane^70^. Thus far, however, little has been elucidated about the structure and mechanism of the eukaryotic CoQ metabolon whose components, the COQ proteins, are all bound to the inner mitochondrial membrane^53, 54^. Here, by combining cryoEM, biochemical experiments, and molecular dynamics simulations, we offer the first structural insight into the actions of a complex of COQ proteins at a lipid bilayer. We present structure of a substrate- and NADH cofactor-bound complex comprised of the hydroxylase COQ7 and the isoprene lipid-binding protein COQ9. Our results reveal how two membrane-bound proteins can cooperate to form a local membrane niche where they can retain, access, and exchange CoQ precursors.

Our data support and extend our previous model that COQ9 facilitates substrate delivery to COQ7^24^; however, the precise mechanism remains elusive. We demonstrate that the COQ7- COQ9 interaction facilitates binding of the CoQ8 precursor OPP, which was enriched 50-fold over background. We further reveal a novel structure^24, 71–73^ of COQ7 with two of its helices forming a membrane-interacting hydrophobic surface and a quinone-binding channel leading to the protein’s active site. However, in the conformation we captured by cryoEM, this channel is not oriented towards COQ9 but towards the interior of the lipid bilayer. It is possible that COQ7 may rotate to accept the substrate from COQ9 as part of an overall cooperative mechanism; however, we did not observe this in our molecular dynamics simulations. Instead, in a separate simulation, we observed that the two membrane-bound helices of COQ7 can open and permit lateral quinone movement. Therefore, we speculate that COQ9 partially desorbs the quinone- intermediate from the membrane and stimulates opening of COQ7’s helices to allow a guided lateral diffusion of the intermediate into COQ7. We further speculate that the tetrameric form of the COQ7:COQ9 complex creates a proteinaceous boundary that protects the intermediates from diffusing away. Interestingly, localization and dynamics of quinones in membranes are known to be influenced by membrane composition and curvature^74, 75^, and the presence of proteins^76^. Therefore, we also hypothesize that the membrane distortion induced by the COQ7:COQ9 tetramer may have an additional function and help to locally retain and concentrate the quinone- intermediates close to the surface of the membrane where they can be efficiently extracted and modified by the enzymes.

A more complete understanding of how the COQ7:COQ9 complex functions will be aided by further establishing if and how its components interact with other COQ proteins. Prior work in yeast and human cells suggests a larger complex Q (or CoQ synthome) that likely includes many or all proteins involved in the processing of CoQ precursors following prenylation by COQ2^8, 9, 12, 14, 50, 63–68^. Gel filtration and 2D blue native-PAGE analyses of digitonin-solubilized yeast mitochondria show that COQ proteins form complexes spanning a wide range of sizes with increased signal detected around 66 kDa, 700 kDa, and 1300 kDa^10, 63, 65–68^. It has been proposed that the 700 kDa complex lacks COQ7 and is dedicated to the production of the DMQ intermediate, whereas the 1300 kDa assembly represents the full complex that finalizes production of CoQ^77^. Although both COQ7 and COQ9 are indeed enriched at the higher molecular weight, they are also detected at the smaller one^10, 67^. Therefore, further studies are required to understand what the complexes of different sizes truly represent. It is possible that complex Q is a singular complex that directly channels quinone-intermediates between its proteins, but that is not sufficiently stable to survive purification. It is also possible that COQ proteins assemble into smaller complexes that co-reside at a specific region of the mitochondrial membrane where they process their substrates. Intriguingly, the stability of COQ proteins and their interactions seems to be positively affected by the presence of CoQ and its intermediates in the membrane^10, 78^. Moreover, fluorescent microscopy studies have shown that in vivo COQ proteins assemble into CoQ-domains and colocalize with ER-mitochondria contact sites in yeast (ERMES)^8, 79^ that facilitate exchange of lipids between the organelles^80^. Whether complex Q and CoQ-domains are one entity, and whether local membrane composition regulates CoQ production remain to be established.

Although overall consistent, our results do have important limitations. Due to the use of a bacterial system of protein expression, we are unable to retrieve demethoxy-coenzyme Q (DMQ), the established substrate of COQ7. Instead, our structures are occupied by octaprenylphenol (OPP), an early intermediate of CoQ8 production in E. coli. Moreover, COQ7 itself appears to be an apo-enzyme, as we do not observe any densities that could represent iron atoms. Unfortunately, a parametrized model of DMQ is not available and iron has very complex coordination chemistry; thus, we are not able to easily overcome these limitations using in silico methods. This prohibits us from making conclusive statements about the catalytic mechanism of COQ7. However, we contend that our structure presents a very plausible quinone-protein arrangement that will guide more detailed studies in the future. Further work will be also required to elucidate what oligomeric forms of the COQ7:COQ9 complex exist in vivo. We suggest the COQ7:COQ9 heterotetramer to be the primary biological unit; however, it cannot be ruled out that other assemblies of COQ7 and COQ9 have biological relevance. It is intriguing that the very stable COQ7:COQ9 octamer is rich in lipids but does not require detergents for purification. How and why two membrane-binding proteins would entrap and solubilize a portion of the plasma membrane is unknown. The membrane curving and thinning observed in the cryoEM and molecular dynamics simulations may indicate full pinching off of a part of the membrane leading to formation of an enzymatic nano-chamber in vivo. Interestingly, COQ7 has been suggested to play additional functions in the cell by participating in mtDNA homeostasis^81, 82^ and stress-signaling^57, 58^. Moreover, COQ7 (called CAT5 in yeast) and YLR202C (ORF partially overlapping with COQ9) were hits in a screen identifying loci important for sterol uptake^83^. Therefore, roles for COQ7 and COQ9 beyond complex Q, and their existence in different oligomeric forms, are possibilities that will require detailed studies.

Overall, our results provide molecular insight into how peripheral membrane proteins can cooperate at a lipid membrane to selectively extract and process an extremely hydrophobic substrate. Although we focused here on CoQ biosynthesis, we anticipate that our findings will have broader utility and contribute to a better understanding of the metabolism of the rich repertoire of hydrophobic molecules residing in and creating cellular membranes.

## Methods

### Site-directed mutagenesis

Yeast Coq7 and Coq9 point mutants were constructed as described in the Q5® Site-Directed Mutagenesis Kit (New England Biolabs) and were confirmed via Sanger sequencing. Plasmid p413 harboring wild type Coq7 was used as a template. In the case of Coq9, pUC19 Coq9-FLAG plasmid was used as a template and successful mutants were cloned into p416 vector using EcoRI and BamHI restriction enzymes and T4 ligase (New England Biolabs). Yeast were transformed as previously described^1^.

To generate pET24a hi6-GB1-Nd38_COQ7 the insert was amplified from pET30 GB1- Nd38_COQ7 with forward primer harboring NdeI cut site and his6 tag and reverse primer harboring EcoRI site. The amplicon and empty pET24a vector were cut with restriction enzymes, ligated with T4 ligase, and confirmed with Sanger sequencing. Bacteria were transformed with standard 30 sec. 42°C heat-shock method.

### COQ7 purification

E.coli Arctic Express cells harboring pET24a hi6-GB1-Nd38_COQ7 (WT or mutants) were grown O/N at 37°C in 25 mL LB media supplemented with geneticin and kanamycin. Cells were refreshed in 3x 2L LB to OD600=0.1 and grew with shaking at 37°C to OD600=0.4. Cultures were chilled to 20°C, 100uM ammonium iron sulfate was added, and protein expression was induced with 100 μM Isopropyl β- d-1-thiogalactopyranoside (IPTG). After the first and the second hour 100 μM ammonium iron sulfate was added again. Protein production was continued for 24h total and then cells were spun 10 min. x 5000g at 4°C. Each 15g of cell paste were resuspended in 35 mL total Lysis Buffer (20 mM HEPES pH 7.5, 200 mM NaCl, 10% glycerol, 5 mM sodium thioglycolate, 1 mM cysteine, 1 μL Benzonase, 5 mM MgCl2, 500 μM phenylmethylsulfonyl fluoride (PMSF), 0.1 mg/mL lysozyme) and incubated at 4°C with slow rotation. Next, the conical tubes were put on ice and the cells were lysed using Bronson sonifier (50% amplitude, 10 sec. ON/60 sec. OFF, 3 cycles). The lysate was spun (10 min. x 50 000g), supernatant was collected and the soluble fraction of COQ7 was subjected to IMAC purification on TALON resin as described later. The pelleted fraction containing membrane-bound COQ7 was resuspended in 10 mL of Lysis Buffer supplemented with 3% digitonin. Sample was loaded into syringe and further homogenized by passing it several times through a needle. The homogenate was incubated at 4°C with rotation for next 2h. Then, it was transferred to oakridge tube and Lysis Buffer was added to 30 mL total. Sample was spun 1h x 50 000g and supernatant containing solubilized proteins was loaded onto 5 mL of TALON resin equilibrated with Wash Buffer (20 mM HEPES 7.5, 200 mM NaCl, 10% glycerol, 1 mM imidazole) in 50 mL conical tube. After 2h incubation at 4°C with gentle rotation, the resin was washed 3 times with 50 mL of Wash Buffer + 10 mM imidazole. Next, the resin was transferred to 15 mL conical tube and the COQ7 was eluted with 5 mL of Elution Buffer (20 mM HEPES 7.5, 200 mM NaCl, 10% glycerol, 150 mM imidazole). Sample was incubated in ice for 10 min., spun 5 min. x 700g, supernatant was collected, and the resin was subjected to another round of elution. Supernatants were combined, concentrated to 1 mL using Amicon Centrifugal Filter (15k Da cut-off) and subjected to Size Exclusion Chromatography. Best fractions were pooled together, frozen with liquid nitrogen and stored at -80C.

### COQ9 and COQ7:COQ9 complex purification

E.coli Arctic Express cells harboring pETduet his6-Nd79_COQ9 alone or co-transformed with pET30 GB1-Nd38_COQ7 were grown O/N at 37°C in 25 mL LB media supplemented with geneticin, ampicillin and kanamycin as needed. Cells were refreshed in 3x 2L LB to OD600=0.1 and grew with shaking at 37°C to OD600=0.4. Cultures were chilled to 20°C, 100 μM ammonium iron sulfate was added, and protein expression was induced with 100 μM Isopropyl β- d-1- thiogalactopyranoside (IPTG). After the first and the second hour 100 μM ammonium iron sulfate was added again. Protein production was continued for 24h total and then cells were spun 10 min. x 5000g at 4°C. Each 15g of cell paste was resuspended in 35 mL total Lysis Buffer (20 mM HEPES pH 7.5, 200 mM NaCl, 10% glycerol, 5 mM sodium thioglycolate, 1 mM cysteine, 1 μL Benzonase, 5 mM MgCl2, 500 μM phenylmethylsulfonyl fluoride (PMSF), 0.1 mg/mL lysozyme) and incubated at 4°C with slow rotation. Next, the conical tubes were put on ice and the cells were lysed using Bronson sonifier (50% amplitude, 10 sec. ON/60 sec. OFF, 3 cycles). The lysate was spun (30 min. x 50 000g) and the supernatant was loaded onto 5 mL of TALON resin equilibrated with Wash Buffer (20 mM HEPES 7.5, 200 mM NaCl, 10% glycerol, 1 mM imidazole) in 50 mL conical tube. After 2h incubation at 4°C with gentle rotation, the resin was washed 3 times with 50 mL of Wash Buffer + 10 mM imidazole. Next, the resin was transferred to 15 mL conical tube and proteins were eluted with 5 mL of Elution Buffer (20 mM HEPES 7.5, 200mM NaCl, 10% glycerol, 150 mM imidazole). Sample was incubated in ice for 10 min., spun 5 min. x 700g, supernatant was collected, and the resin was subjected to another round of elution. Supernatants were combined, concentrated to 1 mL using Amicon Centrifugal Filter (15k Da cut-off) and subjected to Size Exclusion Chromatography and subsequent blue native PAGE analysis (NativePAGE, 4-16%, Bis-Tris, Invitrogen).

### Size Exclusion Chromatography

Purified proteins concentrated to 1 mL or gel filtration standards (BioRad, 1511901) were injected on to Hi-Load pg200 column and 150 mL eluate was collected as 1.0 mL fractions at 1.0 mL/min., in 20 mM HEPES pH 7.5, 200 mM NaCl buffer or in 20 mM ammonium acetate pH 7.0 in case of the COQ7:COQ9 complex subjected later to the cryoEM. UV absorbance signal was used to detect protein-containing fractions which were then analyzed by SDS-PAGE (NuPAGE™ 4 to 12%, Bis-Tris, Invitrogen) to assess their purity. Best fractions were pooled together, frozen with liquid nitrogen and stored at -80°C.

### Lipids extraction from yeast

Yeast were transformed with p413 or p416 plasmids carrying wild type or mutated version of yeast COQ7 and COQ9 genes, respectively. Next, individual colony was picked from the S.C. 2% glucose pABA- plate and used to start 3 mL liquid culture in the same media. After 12h growth at 30°C with shaking, the cultures were used to prepare new 3 mL cultures in S.C. 0.1% glucose, 3.0% glycerol, 50 nM pABA at OD600=0.1 and incubated at 30C with shaking for 24h to pass the diauxic shift. Next, equivalent of 5 mL OD600=1.0 (∼5e7 cells) was pelleted (2 min. x 3000g) in 1.5 mL screw-cap tube. The supernatant was discarded, 100 μL of glass beads, and 50 μL of 150 mM KCl, and 600 μL of methanol with 0.1 μM CoQ8 internal standard (Avanti Polar Lipids) were added, and the cells were lysed by 2 rounds of 5 min. bead-beating in disruptor genie set to max (3000 rpm) speed. To extract lipids, 400 μL of petroleum ether was added to sample and subjected to bead-beating for 3 min. Sample was then spun 2 min. x 1000g at 4°C and the ether layer (top) was transferred to a new tube. Extraction was repeated, the ether layers were combined and dried (∼30 min.) under argon gas at room temperature.

### CoQ6 measurement by HPLC-ECD

Extracted dried lipids were resuspended in 50 μL of mobile phase (78% methanol. 20% isopropanol, 2% 1 M ammonium acetate pH 4.4 in water and transferred to amber glass vials with inserts. Vials were inserted into HPLC (Ultimate 3000, Thermo Scientific) with electrochemical detector (ECD-3000RS) and 10 μL of each sample was injected. The flow rate was set to 0.3 mL/min (LPG-3400RS pump). The first electrode (6020RS) was set to +600 mV and placed before the column (Thermo Scientific, Betasil C18, 100 x 2.1 mm, 3um particle) to oxidize all the quinones and ensure their simultaneous elution. Second electrode (6011RS) was set to -600 mV to reduce the quinones exiting the column, and then the third electrode was set to +600 mV to make final recordings. A CoQ6 standard (Avanti Polar Lipids) was used to identify corresponding peak in the obtained data. Peaks were then quantified with Chromeleon 7.2.10 software using cobra wizard option.

### Measurement of DMQ0’s redox state by HPLC-ECD

Samples were prepared according to “NADH fluorescence method”. After 30 min. incubation 50 μL of each sample was quenched with 50 μL of -20°C cold methanol, incubated at -20°C for 5 min. and spun at 2°C for 5 min. x 20 000 g. Supernatants from triplicates were pooled together and transferred to amber glass vials. Vials were inserted into HPLC (Ultimate 3000, Thermo Scientific) with electrochemical detector (ECD-3000RS) and cooled to 4°C. Then, 2.5 μL of each sample was injected and resolved at 0.25 mL/min. (LPG-3400RS pump) isocratic flow of 90% A (0.1% formic acid in H2O) and 10% B (isopropanol) mobile phase. The first electrode (6020RS) placed before the column (BetaBasic C18, 100 x 4.5 mm, 5um particle) was turned off (0 mV). Second electrode (6011RS) was set to -600 mV to reduce the quinones exiting the column, and then the third electrode was set to +600 mV to make final recordings. Data were compared to 250 μM DMQ0 standard (WuXi AppTec) pre-oxidized with first electrode set to +600 mV or pre-reduced with the electrode set to -600 mV.

### Wester Blotting

Methanol-glass beads mix from the lipid extraction protocol was dried under fume hood at 60°C for several hours until complete dryness. Next, 200 μL of 1x LDS Buffer with 10 mM DTT was added and the sample was boiled for 20 min. with often intense vortexing. Beads were spun 5 min. x 20 000g and 15 μL of supernatant was loaded on SDS-PAGE gel (NuPAGE™ 4 to 12%, Bis-Tris, Invitrogen). Proteins were electrophoretically separated for 35 min. at 150V, and then transferred (192 mM glycine, 25 mM Tris, 20% methanol [v/v]) to methanol-activated PVDF membrane (Immobilon-FL). The membrane was washed with TBS-T (20 mM Tris pH 7.4, 150 mM NaCl, 0.05% Tween 20 [v/v]), blocked with 5% powdered milk in TBS-T, washed 3 times with TBS-T, incubated at 4°C O/N with primary α-FLAG (Millipore F1804-5MG) or α- beta-Actin (Abcam ab8224) or α-Coq7 (raised by GenScript in rabbits against NLERTDGTKGPSEE peptide) antibodies (1:5000 in 0.5% milk), washed 3 times with TBS-T, incubated with secondary antibodies (1:20 000 in 0.1% milk, goat anti-mouse (LI-COR 926-32210, 1:15000; RRID: AB_621842) or goat anti-rabbit (LI-COR 926-32211, 1:15000; RRID: AB_621843)), washed 3 times with TBS-T, and finally visualized with LI-COR Odyessey CLx scanner using Image Studio v5.2 software.

### Yeast respiratory growth

Individual colony was picked from the S.C. -URA or -HIS, 2% glucose, pABA-, plate and used to start 3 mL liquid culture in the same media. After 12h growth at 30°C with shaking, the culture was used to prepare new 1.0 mL culture in S.C. 0.1% glucose, 3.0% glycerol, 50 nM pABA at OD600=0.05. Next, 100 μL of it was transferred to 96-well round-bottom clear plate (Thermo) and sealed with BreatheEasy film (USA Scientific). The plate was loaded into EPOCH plate reader operated with Gen5 3.1 software and yeast were grown at 30°C with max linear shaking (1096 cpm) for 48-72h with optical density monitored every 20 min. at 600 nm.

### COQ7:COQ9 complex delipidation

100 μL of COQ9:COQ7 complex was thawed (20mM HEPES pH 7.5 200mM NaCl, 10% glycerol at 1.5 μg/μL) and incubated 1h with 100 μL BioBeads (washed with 1mL of methanol and then 3x 1mL buffer) at room temp with orbital shaking. In parallel, 100 μL of COQ9:COQ7 was incubated without biobeads as a control of protein stability at the room temp. 5 μL was collected after 1h. The beads were spun 15 000g x 1 min. and the supernatant containing delipidated complex was transferred to a new tube. The beads were washed 3 times with 1 mL of the buffer and resuspended in 100 μL of 1x LDS buffer with 5 mM DTT. Finally, blue native PAGE (NativePAGE, 4-16%, Bis-Tris, Invitrogen) and SDS-PAGE (NuPAGE™ 4 to 12%, Bis-Tris, Invitrogen) were run to analyze results.

### NADH fluorescence

Buffer (20 mM HEPES pH 7.6, 200 mM NaCl) was mixed with 5 μM protein, 250 μM NADH, and 250 μM DMQ0 as indicated in the figure to total reaction volume of 100 μL. Samples were loaded into black flat clear bottom 96-well plate (BRANDplates, pure-grade) and incubated at 30°C with constant slow orbital shaking. Optics were set to TOP position with probe’s height at 5 mm and gain value 100. Then, NADH’s fluorescence was recorded every 5 min. (Excitation= 340 nm, Emission=445 nm) for 4h.

### Differential Fluorimetry Scanning (DSF)

24.5 μL of 5 μM solution of every protein was prepared in 96-well plate MicroAmp Optical plate (Applied biosystems), and incubated at room temp. for 15 min. Then, 0.5 μL 50x SYPRO Orange (Invitrogen) dye in 10% DMSO was added to each well and mixed by pipetting. The plate was sealed with MicroAmp Optical film (Applied biosystems), spun 1 min. x 1000 g, loaded into QuantStudio 6 Flex (Applied biosystems) and subjected to 20-99°C heat gradient changing at 0.025 °C/sec. Signal for ROX dye (overlapping with SYPRO orange) was recorded. The data were analyzed using DSFworld server (https://gestwickilab.shinyapps.io/dsfworld). Fits 2 and 4 were selected as the best representing the shape of melting curves and used to calculate melting temperatures.

### In silico analyses

Tunnels in COQ7 were detected with MOLEonline^2^ server (https://mole.upol.cz/) using default settings. All graphs were prepared using GraphPad Prism 9.1.0 software. All structures were analyzed (inter-residue distances, interactions etc.) and visualized using CHIMERA 1.11.2 software^3^. Electrostatic surface potentials of proteins were calculated using APBS server^4^ (poissonboltzmann.org) on default settings. Densitometry was performed with ImageJ 1.50i^5^. Student t-tests were calculated in Excel.

### Mass-spec lipidomics analysis of DMQ6 content in yeast

Lipid extracts from equivalents of 5 mL OD600=1.0 (∼5e7 cells) yeast cultures were prepared as described in “HPLC-ECD” section. LC-MS analysis was performed using Thermo Vanquish Horizon UHPLC system coupled to a Thermo Exploris 240 Orbitrap mass spectrometer. For LC separation, A Vanquish binary pump system (Thermo Scientific) was used with a Waters Acquity CSH C18 column (100 mm × 2.1 mm, 1.7 mm particle size) held at 35°C under 300 μL/min flow rate. Mobile phase A consisted of 5 mM ammonium acetate in ACN/H2O (70:30, v/v) containing 125 μL/L acetic acid. Mobile phase B consisted of 5 mM ammonium acetate in IPA/ACN (90:10, v/v) with the same additive. For each sample run, mobile phase B was initially held at 2% for 2 min and then increased to 30% over 3 min. Mobile phase B was further increased to 50% over 1 min and 85% over 14 min and then raised to 99% over 1 min and held for 4 min. The column was re-equilibrated for 5 min at 2%B before the next injection. Five microliters of sample were injected by a Vanquish Split Sampler HT autosampler (Thermo Scientific) while the autosampler temperature was kept at 4 °C. The samples were ionized by a heated ESI source kept at a vaporizer temperature of 350°C. Sheath gas was set to 50 units, auxiliary gas to 8 units, sweep gas to 1 unit, and the spray voltage was set to 3,500 V for positive mode and 2,500 V for negative mode (two separate injections per replicate). The inlet ion transfer tube temperature was kept at 325°C with 70% RF lens. For discovery analysis, MS^1^ scans were acquired at 120,000 resolution from m/z 200 to 1,700 with EasyIC enabled to improve mass accuracy. MS^2^ scans were acquired in AcquireX Background Exclusion mode to automatically exclude background ions found in the extraction blank from MS/MS fragmentation during sample acquisitions with the cycle time of 1.5 s. Other MS^2^ parameters include resolution of 30,000, 54 ms MS^2^ ion injection time, 1.5 m/z isolation width, stepped HCD collision energy (25%, 30% for positive mode and 20%, 40%, 60% for negative mode), and 3 s dynamic exclusion. Automatic gain control (AGC) targets were set to standard mode for both MS^1^ and MS^2^ acquisitions.

### Data Analysis

The resulting CoQ intermediate data were processed using TraceFinder 5.1 (Thermo Fisher Scientific).

### LC-MS lipidomics of purified proteins and E. coli cells

For lipidomics analysis of COQ7:COQ9 complex, two 20 uL aliquots of 5 mg/mL protein COQ9:COQ7 complex, in 200 mM ammonium Acetate were extracted using Matyash method^6^. Briefly, once samples were thawed on ice, 200 μL methyl tert-butyl ether (MTBE), 60 uL of methanol, and 50 uL of water was added to each tube and the mixture was vortexed for 10 s. The sample was then centrifuged for 10 min at 10,000 g at 4°C. 150 μL of the lipophilic (upper) layer from the biphasic extraction was aliquoted into a separate glass vial and dried down by vacuum centrifugation, and was then resuspended in 50 μL 9:1 MeOH/Toluene prior to LC-MS/MS analysis.

For lipidomics of the separately purified proteins, two aliquots of each purified protein (100 μg COQ7, 100 μg COQ9, 200 μg COQ7:COQ9) or pelleted E.coli cells (10e10, ∼20 x OD600=1.0) were thawed on ice. Once thawed, 87 μL of methanol, and 290 μL methyl tert-butyl ether (MTBE) were added to each non-pellet tube and the mixture was vortexed for 10 s (sample contained 100 μL of aqueous buffer). For each of the E. coli cell pellet tubes, 205 μL methanol, 750 μL MTBE, and 187.5 μL water were added, and the mixture was sonicated for 10 s. After centrifugation for 2 min at 14,000 g at 4°C, 200 μL of the lipophilic (upper) layer from the biphasic extraction was aliquoted into an amber glass autosampler vial with glass insert and dried down by vacuum centrifugation. The samples were then resuspended in 50 μL 9:1 MeOH/toluene prior to LC-MS/MS analysis.

### LC-MS/MS analysis for lipidomics

Sample analysis was performed using an Acquity CSH C18 column held at 50 °C (100 mm x 2.1 mm x 1.7 μm particle size; Waters), a 400 μL/min flow rate was maintained using a Vanquish Binary Pump (Thermo Scientific). The mobile phases consisted of 10 mM ammonium acetate in ACN:H2O (70:30, v/v) with 250 μL/L acetic acid (Mobile Phase A), and 10 mM ammonium acetate in IPA:ACN (90:10, v/v) with 250 μL/L acetic acid (Mobile Phase B). For each sample 10 μL, was injected onto column using Vanquish autosampler (Thermo Scientific). The gradient for lipid separations was as follows, mobile phase B was initially held at 2% for 2 min and then increased to 30% over 3 min. Mobile phase B was further increased to 50% over 1 min, then raised to 85% over 14 min, and finally raised to 99% over 1 min and held at 99 % for 7 min. The column was re-equilibrated with mobile phase B at 2% for 1.75 min before the next injection.

The LC system was coupled to a Q Exactive Orbitrap mass spectrometer through a heated electrospray ionization (HESI II) source (Thermo Scientific). Source conditions were as follow: HESI II and capillary temperature at 300 °C, sheath gas flow rate at 25 units, aux gas flow rate at 15 units, sweep gas flow rate at 5 units, spray voltage at |3.5 kV| for both positive and negative modes, and S-lens RF at 90.0 units. Data were acquired using polarity switching with positive and negative full MS and MS2 spectra (Top2) within the same injection. Acquisition parameters for full MS scans in both modes were 17,500 resolution, 1 × 10^6^ automatic gain control (AGC) target, 100 ms max inject time, and 200 to 1600 m/z scan range. MS2 scans in both modes were then performed at 17,500 resolution, 1 × 10^5^ AGC target, 50 ms max IT, 1.0 m/z isolation window, stepped normalized collision energy (NCE) at 20, 30, 40, and a 10.0 s dynamic exclusion

### Data Analysis for lipidomics by mass spectrometry

LC–MS data were processed using Compound Discoverer 2.1 and 3.1 (Thermo Scientific) and LipiDex, an in-house-developed software suite^7^. Features detection was performed within a 1.4 min to 21 min retention time window, and features were aggregated into compound groups using a 10-ppm mass and 0.75 min retention time tolerance. Detect compounds node settings of minimum peak intensity of 5×10^5^, maximum peak-width of 0.75, and signal-to-noise (S/N) ratio of 3 were used. Background features were designated as features less than 3-fold intensity over blanks and were excluded from further processing. MS/MS spectra were searched against an in-silico generated lipid spectral library^7^. Spectral matches with a dot product score greater than 500 and a reverse dot product score greater than 700, eluting within a 3.5 median absolute retention time deviation (M.A.D. RT) of each other, and found within at least 2 processed files were retained. Lipids with no significant interference (<75 %) from co-eluting isobaric lipids were identified at the individual fatty acid substituent level, otherwise lipids were annotated with the sum of the fatty acid substituents.

### In-gel sample digestion for proteomics

All the steps were performed at room temperature with use of LC-MS grade reagents. Samples were loaded onto reducing SDS-PAGE gel (NuPAGE™ 4 to 12%, Bis-Tris, Invitrogen) and resolved 45 min. x 150V. Individual bands were excised from the gel, placed in separate tubes and cut into smaller pieces. Samples were destined for 15 min. in 100 μL of 25 mM ammonium bicarbonate (AMBIC) in acetonitrile/water (50/50, v/v) buffer. Next, samples were washed for 10 min. with acetonitrile and air dried for 10 min. Then, samples were reduced (40 μL of 10 mM TCEP and 40 mM 2-chloroacetemide in 25 mM AMBIC) for 30 min. and dehydrated in 200 μL acetonitrile for 10 min. The gel pieces were air dried for 10 min., incubated with 40 μL of trypsin (Promega, Sequencing Grade, 16 ng/μL in 25 mM AMBIC) for 15 min., supplemented with additional 30 μL of 25 mM AMBIC and the proteins were digested overnight. Supernatants containing digested peptides were transferred to new tubes and gel pieces were covered with 100 μL extraction buffer (5% formic acid in water/acetonitrile (1/2, v/v), incubated for 10 min. and the supernatants were collected and combined with the previous ones. Samples were dried down to several μL in Speedvac (Thermo Scientific), resuspended in 100 uL 0.1% TFA and desalted using C18 Omix tips (Agilent, #A57003100K). Each time a tip was conditioned with 2x 100 μL of 0.1% TFA in 50:50 (v/v) ACN/H2O, washed with 2x 100 μL of 0.1% TFA in H2O, sample was bound by passing 10 times thru tip resin, and then washed 3x 100 μL of 0.1% TFA in H2O.

The peptides were eluted with 100 μL of 0.1% TFA in 80:20 (v/v) ACN/H2O, dried in Speedvac, reconstituted in 20 μL of 0.2% formic acid in H2O and subjected to LC-MS analysis.

### Liquid Chromatography Mass Spectrometry Proteomics

LC separation was performed using Thermo Ultimate 3000 RSLCnano system. A 15 cm EASY- Spray™ PepMap™ RSLC C18 column (150 mm × 75 μm, 3 μm) was used at 300 nL/min flow rate with a 60 min. gradient using mobile phase A consisting of 0.1% formic acid in H2O, and mobile phase B consisting of 0.1% formic acid in ACN/H2O(80/20, v/v). EASY-Spray source was used and temperature was at 35 °C. Each sample run was held at 4.0% B for 3 min and increased to 45% B over 42 min, followed by 5 min at 95% B and back to 4% B for equilibration for 10 min. An Acclaim PepMap C18 HPLC trap column (20 mm × 75 μm, 3 μm) was used for sample loading. MS detection was performed with Thermo Exploris 240 Orbitrap mass spectrometer in positive mode. The source voltage was set to 1.5 kV, ion transfer tube temperature was set to 275 °C, RF lens was at 40%. Full MS spectra were acquired from m/z 350 to 1400 at the Orbitrap resolution of 120000, with the normalized AGC target of 300% (3E6) and max ion injection time of 25 ms. Data-dependent acquisition (DDA) was performed for the top 15 precursor ions with the charge state of 2-6 and an isolated width of 2. Intensity threshold was 5E3. Dynamic exclusion was 20 s with the exclusion of isotopes. Other settings for DDA include Orbitrap resolution of 15000, HCD collision energy of 30%, and ion injection time of 40 ms.

Raw data files were analyzed by Andromeda Search engine incorporated in MaxQuant v1.6.17.0 software against human and E. Coli databases downloaded from Uniprot. Label-free quantitation was enabled in the analysis to compute iBAQ values for the identified proteins.

### Negative-stain electron microscopy

4.0 µl aliquots of purified COQ7-COQ9 complex (0.05 mg ml^-1^) was applied onto glow-discharged continuous carbon-coated grids, blotted with filter paper, and stained with freshly prepared 0.75% (w/v) uranyl formate. Negatively-stained electron microscopy grids were imaged on Tecnai T12 microscope (FEI Company) equipped with a 4k x 4k CCD camera (UltraScan 4000, Gatan) and operated at a voltage of 120 kV. Images were recorded at room temperature with a nominal magnification of 52,000x, corresponding to a calibrated pixel size of 2.21 Å on the specimen. A total of 121 micrographs were collected and CTF parameters were estimated by CTFFIND4^8^. Particles were manually picked, extracted with a box size of 200 x 200 pixels, and reference-free 2D classification was used to generate the templated for the automatic particle picking by using Relion 3.0 software package^9^. Then, 262,589 particle projections were semi-automatically picked from the micrographs and sorted through multiple rounds of 2D classification to discard poorly defined particle classes using Relion 3.0. A subset of 231,879 particles with well-defined shapes was selected following multiple rounds of 2D classification to demonstrate 2D class averages of COQ7-COQ9 complex.

### Cryo-EM sample preparation

Peak fractions containing the COQ7-COQ9 complex were concentrated to 5 mg ml^-1^ and flash frozen in liquid nitrogen for long-term storage at -80 °C. An aliquot of purified COQ7-COQ9 complex was diluted to 0.5 mg ml^-1^ (8.8 µM) in 200mM Ammonium Acetate, pH 7.0 before blotting. Cryo-EM samples of COQ7-COQ9 complex bound to NADH were prepared similarly with the addition of NADH. Briefly, the diluted complex (8.8 µM) was mixed with NADH to a final NADH concentration of 5 mM in a final solution containing 200mM Ammonium Acetate, pH 7.0, and incubated at room temperature for 1 hour. To prepare cryo-EM grids, 3.5 µl of purified COQ7-COQ9 complex was applied to glow-discharged holey carbon grids (Quantifoil R1.2/1.3, 400 mesh Cu) and incubated for an additional 30 s. Then, the grids were blotted with Whatman Grade 1 filter paper (Whatman) for ∼6 s with a 0 mm offset at 10 °C and 100% relative humidity and plunge frozen into liquid ethane using a Vitrobot Mark IV (Thermo Fisher Scientific).

### Data collection

Two datasets were collected on two different microscopes using the multi-record strategy (nine-hole exposures per single-stage movement) in SerialEM software^10^. For unliganded COQ7-COQ9 complex, images were collected on a FEI Talos Arctica equipped with a K3 Summit direct electron detector (Gatan) and operated at an accelerating voltage of 200 keV. In total, 1,395 images were recorded with a defocus range of -0.5 to -1.5 µm in super-resolution counting mode at a nominal magnification of 36,000x, corresponding to a super-resolution pixel size of 0.57 Å (physical pixel size of 1.14 Å) on the specimen level. Each image was dose-fractionated over 120 frames with a per frame dose rate of ∼0.49 electrons and total exposure time of 2.4 s, resulting in an accumulated dose of ∼58.8 electrons per Å^2^. For NADH-bound COQ7-COQ9 complex, images were acquired on a FEI Titan Krios equipped with a K3 Summit direct electron detector and a Quantum GIF energy filter (Gatan) with a slit width of 20 eV and operated an accelerating voltage of 300 keV in nano-probe mode. A total of 10,088 micrographs were collected at a magnification of 105,000x with a calibrated super-resolution pixel size of 0.4165 Å (physical pixel size of 0.833 Å) and a defocus range of -0.3 to -1.2 µm. The total exposure time for each micrograph was 3 s, with dose-fractionation set at 0.025 s per frame, resulting in 118 movie frames at a per frame dose rate of ∼0.55 electrons for a total accumulated dose of ∼65 electrons per Å^2^. Data collection statistics are shown in Extended Data Table 1.

### Image analysis and 3D reconstruction

Dose-fractionated stacks were subjected to beam-induced motion correction using MotionCor2^11^. CTF parameters for each micrograph in the unliganded COQ7-COQ9 complex and the NADH-bound COQ7-COQ9 complex datasets were determined by GCTF v.1.06^12^ and CTFFIND4^8^, respectively. Subsequent image processing for both datasets was carried out in Relion 3.0. For cofactor-free COQ7-COQ9 complex, 3040 particles were manually picked and classified by reference-free 2D classification to generate templates for automatic particle picking. A total of 574,760 auto-picked particles were extracted with a box size of 192 pixels and subjected to two rounds of 2D classification to discard poorly defined classes, resulting in 528,971 particles for further processing. An initial 3D model from these particles was generated by using cryoSPARC ab initio reconstruction^13^. Stable classes were then used for iterative rounds of 3D refinement and reclassification without symmetry. Upon visual inspection of 3D reconstructions in UCSF Chimera^3^, the particles from the best 3D classes (281,866 particles) were combined and subjected to 3D refinement with D2 symmetry. Subsequent per-particle CTF refinement and 3D classification without alignment yielded a single class containing 61,526 particles. This final subset of particles was refined with a soft mask and sharpened by a negative temperature factor to a resolution of 3.5 Å during post-processing procedure. A similar strategy was used for NADH-bound COQ7-COQ9 complex. In total, 2,709,398 particles were extracted with a box size of 256 pixels and binned to 128 pixels. After removing the suboptimal particles with two rounds of reference-free 2D classification, the best classes were subjected to 3D refinement without imposed symmetry. Subsequent 3D classification into 6 classes showed two dominant classes containing 1,249,119 particles (∼46% of the dataset), which was then re-extracted to a box size of 256 pixels and refined with D2 symmetry imposed. These particles were subjected to another round of 3D classification into six classes without image alignment, resulting in 372,917 particles with indicated global resolution of 2.7 Å. The map was further improved after CTF and aberration refinements by using Relion 3.1^9^. The final map had a resolution of 2.4 Å after sharpened by applying a negative temperature factor during the post-processing step. The reported final resolution estimates are based on the gold-standard Fourier shell correlation (FSC) cut-off of 0.143. Local resolutions were determined using ResMap with half-reconstructions as input maps. Refinement parameters for the final density maps are listed in Extended Data Table 1.

### Model building and validation

The initial model of COQ7 was a homology model calculated by Phyre2^14^, using the intensive modeling mode. The crystal structure of COQ9 (PDB ID: 6AWL) was also used as an initial template for model building. The atomic models were manually rebuilt using Coot^15^ and real-space refined using phenix.real_space_refine tool^16^ from the PHENIX software package^17^. Briefly, phenix.mtriage^17^ was used to analyze cryo-EM maps and an automated sharpening procedure (phenix.auto_sharpen^10^) was applied to the final cryo-EM reconstructions prior to model building. The 3.7 Å and 2.4 Å density maps were of sufficient quality for de novo model building. After initial model building, the unliganded and NADH-bound starting models were subjected to iterative rounds of refinement in phenix.real_space_refine using global minimization, morphing, noncrystallographic symmetry (NCS), B-factor refinement, local grid search and secondary structure restraints, and manual rebuilding in Coot. The ligand coordinates were docked into densities and refined using Coot. The final model geometry was evaluated using MolProbity 4.5^18^. Independent FSC curves for model-map correlations were calculated between the resulting model and the half map used for refinement as well as between the resulting model and the other half map for cross-validation. Protein interfaces and associated free energies were analyzed by using PDBePISA server^19^. The final refinement statistics are summarized in Extended Data Table 1.

### Molecular dynamics simulations

We prepared all systems for MD simulations with the membrane module of the CHARMM-GUI server^20^ and followed the provided equilibration and production files, except for changes as indicated. We parametrized the protein components with the CHARMM36m forcefield, its adapted TIP3P water model and the CHARMM36 lipids^21^, plus the NADH parameters from^22^ and the CoQ10 parameters from^23, 24^. The model membranes contained 17% cardiolipin (charged -2), 44% POPC and 39% POPE; or 16% cardiolipin, 42% POPC and 38% POPE when 4% CoQ10 was included. The solvent included 0.15 M KCl for charge neutralization and ionic strength. For simulations aimed at testing unbinding of CoQ10 from COQ7’s active site, we first tried replica exchange MD but it invariably resulted in membrane destabilization. Therefore, we took another approach in which we ran regular simulations at 300, 400, 450, 500 and 600 K. At 500 and 600 K, the system destabilized within the first few hundred nanoseconds of simulation times: the proteins unfolded and the membranes disrupted. At 300 and 400 K, the small molecule remained locked inside the active site for the full length of independent multi-microsecond trajectories. At 450 K we observed events in which the docked CoQ10 dissociated from the active site without compromising the protein or membrane stabilities within the simulated timescale (∼1-1.5 microsecond); we therefore analyzed 10 independent replicas at this temperature as described in the main text. In all these simulations the membranes and proteins remained stable, and the NADH remained stably bound to COQ7, during the whole simulation times.

We ran all simulations with Gromacs 2020^25^, visualized them with VMD^26^ and analyzed them with custom scripts, VMD procedures, and the MEMBPLUGIN plugin for VMD^27^.

### Data and Code Availability

The raw LC-MS lipidopmics data generated during the current study are available from the corresponding author on reasonable request.

The atomic coordinates of NADH-bound COQ7:COQ9 complex and COQ7:COQ9 complex only have been deposited in the Protein Data Bank with accession numbers 7SSS and 7SSP, respectively. All of the 3D cryoEM density maps associated with this study have been deposited in the Electron Microscopy Data Bank under accession numbers EMD-25413 for NADH-bound COQ7:COQ9 complex and EMD-25412 for COQ7:COQ9 complex only.

No custom computer code was generated for this project.

## Acknowledgments

We thank current and former members of the D.J.P. and A.F. laboratories for helpful discussions and assistance on this project. We specially thank Danielle C. Lohman for pioneering the purification of the COQ7:COQ9 complex that paved the way for its experimental characterization. We thank Annie Jen and Zixiang Fang for their help with LC-MS measurements. We also thank Professor Stephen J. Lippard for kindly providing the pET30 GB1-Nd38_COQ7 vector. M. Braunfeld, D. Bulkley, M. Harrington, A. Myasnikov, and Z. Yu of the UCSF Center for Advanced CryoEM for microscopy support and J. Baker-LePain and the QB3 shared cluster (NIH grants 1S10OD021596-01, S10OD020054, S10OD026881, 1S10OD021741 and the Howard Hughes Medical Institute).

This work was supported by NIH awards R35 GM131795 (D.J.P.), P41 GM108538 (J.J.C. and D.J.P.), support from the Howard Hughes Medical Institute Faculty Scholar program and the Chan Zuckerberg Biohub (A.F.); funds from the BJC Investigator Program (D.J.P.); SNSF grants 205321_192371 and 31003A_170154 (M.D.P). H.A. is a Human Frontier Science Program Long-Term Fellow supported by The International Human Frontier Science Program Organization (LT000398/2017-L).

## Author contributions

**Mateusz Manicki:** Conceptualization, Methodology, Formal analysis, Investigation, Writing – original draft, Writing – review and editing, Visualization, Project Administration. **Halil Aydin:** Conceptualization, Methodology, Software, Formal analysis, Investigation, Data curation, Writing – review and editing. **Luciano A. Abriata:** Conceptualization, Methodology, Software, Formal analysis, Investigation, Data curation, Writing – review and editing, Visualization. **Katherine A. Overmyer:** Methodology, Formal analysis, Investigation, Data curation. **Rachel M. Guerra:** Conceptualization, Methodology, Writing – review and editing. **Joshua J. Coon:** Supervision, Funding acquisition. **Matteo Dal Peraro:** Methodology, Writing – review and editing, Supervision, Funding acquisition. **Adam Frost:** Conceptualization, Methodology, Writing – review and editing, Supervision, Funding acquisition. **David J. Pagliarini:** Conceptualization, Methodology, Writing – review and editing, Supervision, Project administration, Funding acquisition.

## Correspondence and requests for materials

Requests should be addressed to A.F. or D.J.P.

---

**Extended Data Figure 1.**
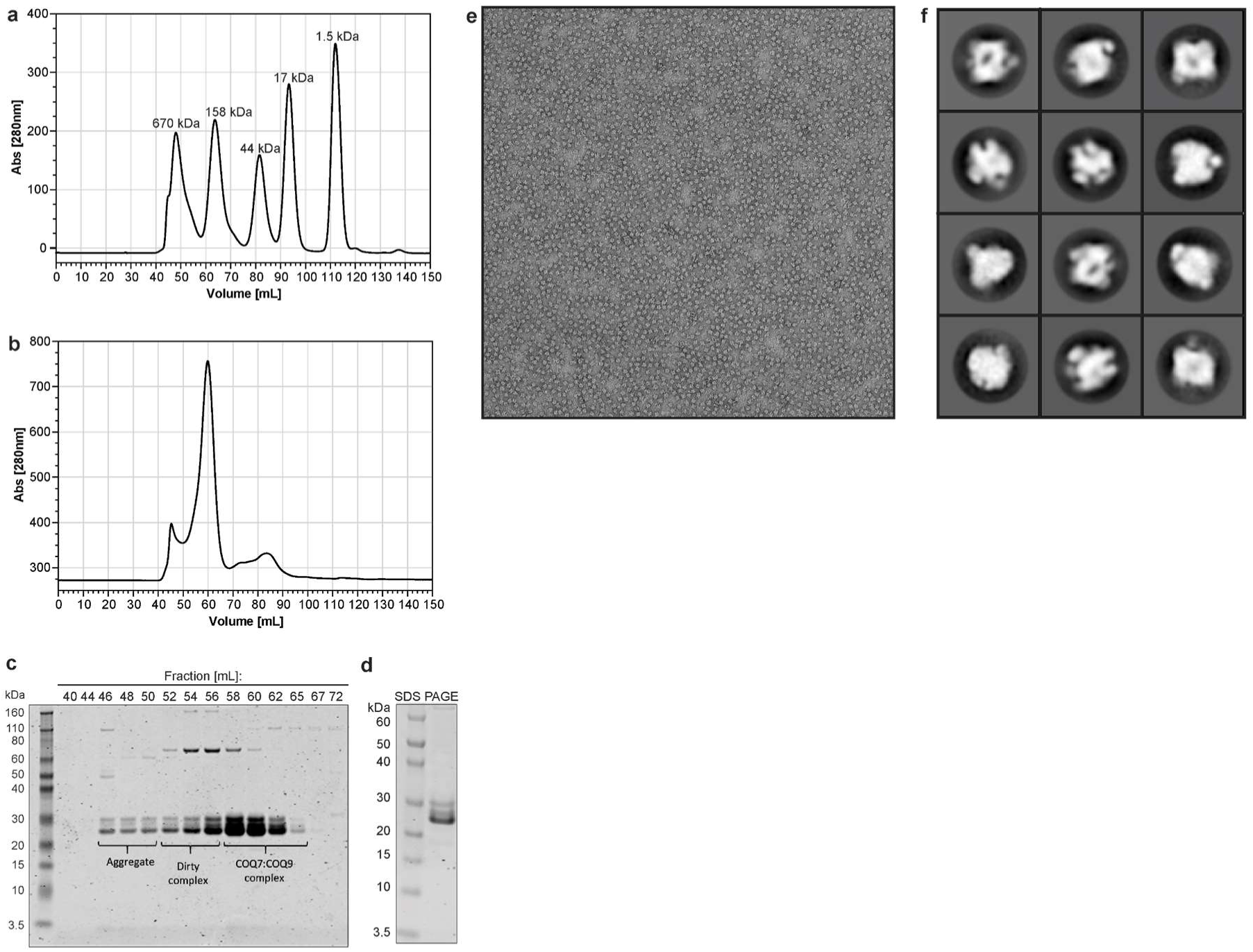
Preliminary biochemical and structural analysis of the COQ7:COQ9 complex. **a**, Size exclusion chromatography trace of protein mass standards. **b**, Size exclusion chromatography trace of the purified GB1-^Nd38^COQ7:His6-^Nd79^COQ9 complex **c**, SDS-PAGE analysis of collected fractions. **d**, SDS-PAGE analysis of pooled GB1-^Nd38^COQ7:His6-^Nd79^COQ9 fractions submitted to cryoEM analysis. **e**, Representative negative-stain EM image of COQ7:COQ9 complex reveals mono-disperse particle distribution with very little protein aggregation. **f**, Representative 2D class averages of COQ7:COQ9 complex from the negative-stain data collected on Tecnai T12 microscope equipped with CCD camera showing that particles have well-defined shapes and symmetric features.

**Extended Data Figure 2.**
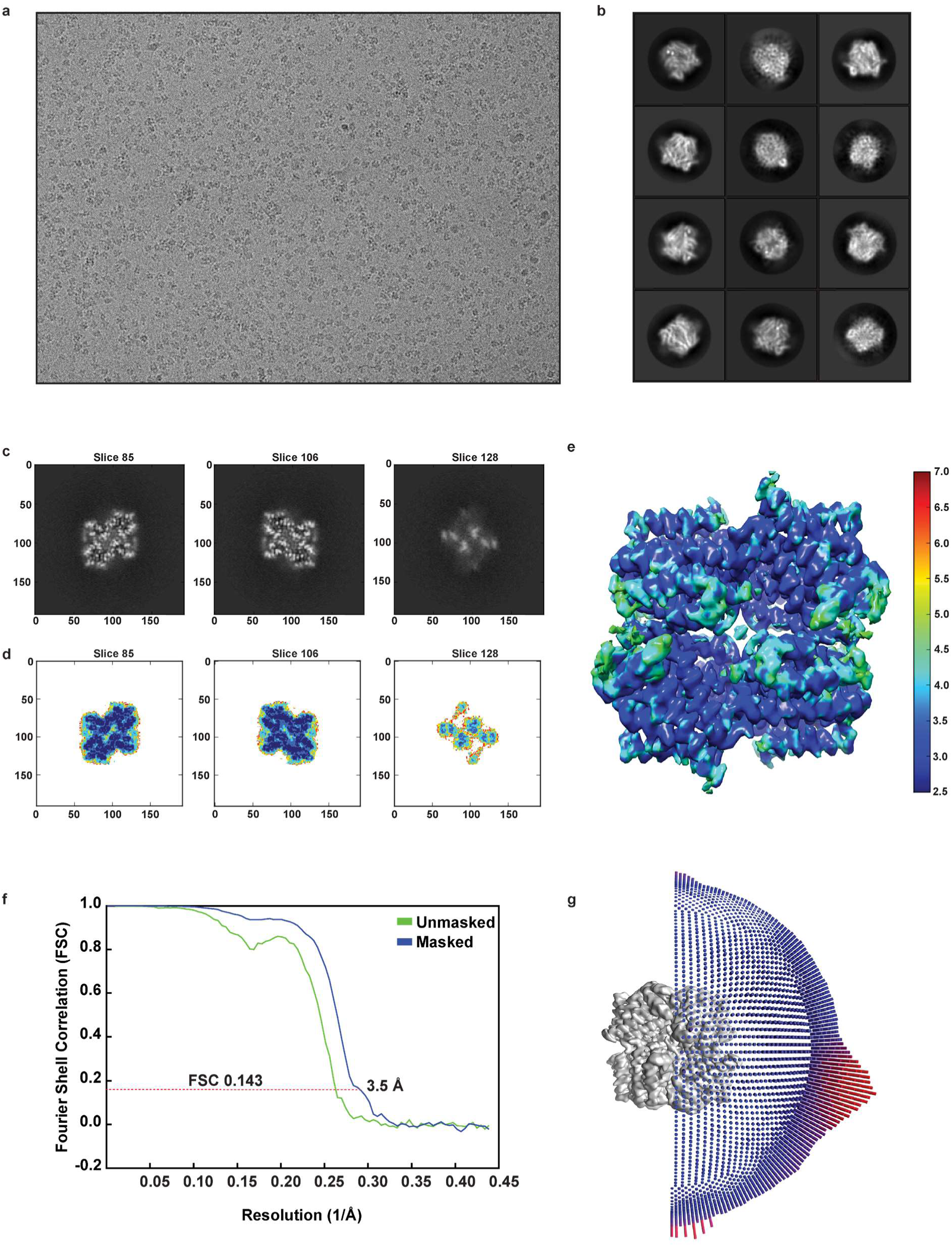
Single-particle cryoEM of unliganded COQ7:COQ9 complex. **a**, Representative motion-corrected electron micrograph of COQ7:COQ9 complex only. **b**, Gallery of 2D class averages calculated from the cryoEM data of the complex. **c, d**, Slices through the unsharpened density map at distinct levels are shown in top view. **e**, Final 3D electron density map colored according to local resolution is shown in side view. Local resolution was calculated by ResMap. **f**, Fourier shell coefficient (FSC) curves (cutoff of 0.143) between two independently refined half maps before and after post-processing. **g**, Orientation distribution of all particle images included in the calculation of the final 3D reconstruction.

**Extended Data Figure 3.**
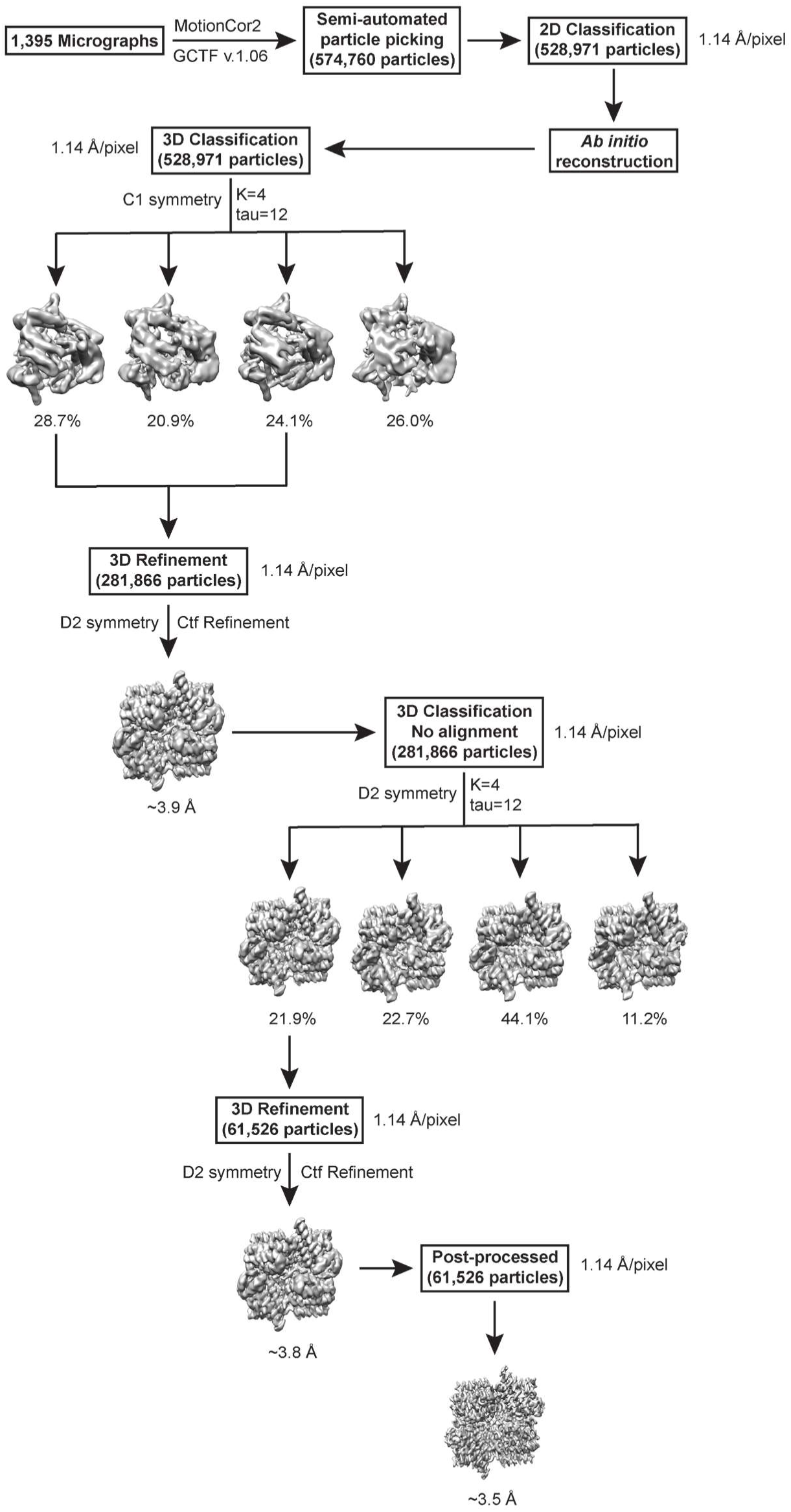
Stages of preprocessing, image classification, refinement, and postprocessing for unliganded COQ7:COQ9 complex. Details can be found in the image analysis and 3D reconstruction section of the Methods.

**Extended Data Figure 4.**
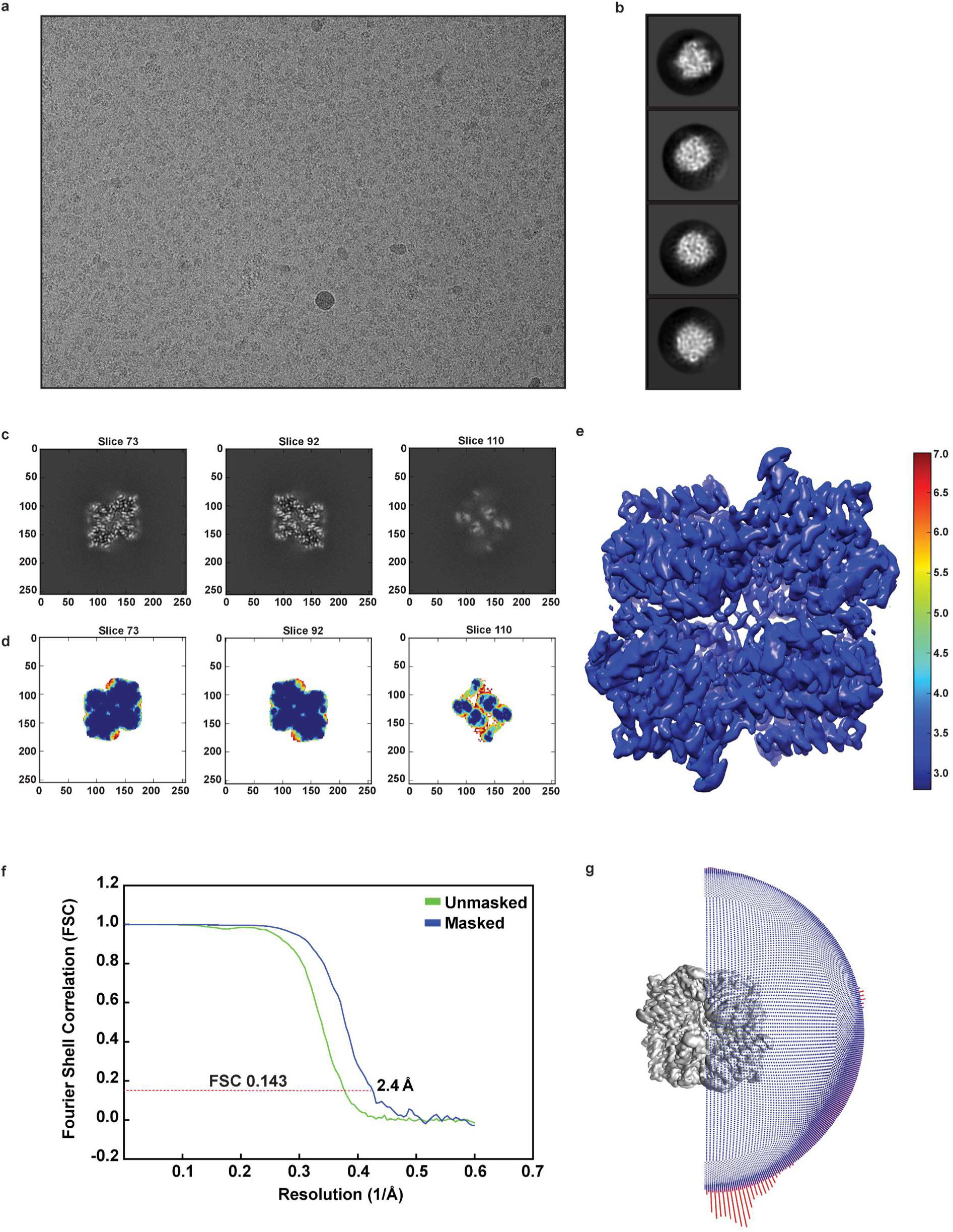
Single-particle cryoEM of NADH-bound COQ7:COQ9 complex. **a**, Representative motion-corrected electron micrograph of NADH-bound COQ7:COQ9 complex. **b**, Representative 2D class averages calculated from the cryoEM data of the complex. **c,d**, Slices through the unsharpened density map at distinct levels are shown in top view. **e**, Final 3D electron density map colored according to local resolution is shown in side view. Local resolution was calculated by ResMap. **f**, Fourier shell coefficient (FSC) curves (cutoff of 0.143) between two independently refined half maps before and after post-processing. **g**, Orientation distribution of all particle images included in the calculation of the final 3D reconstruction.

**Extended Data Figure 5.**
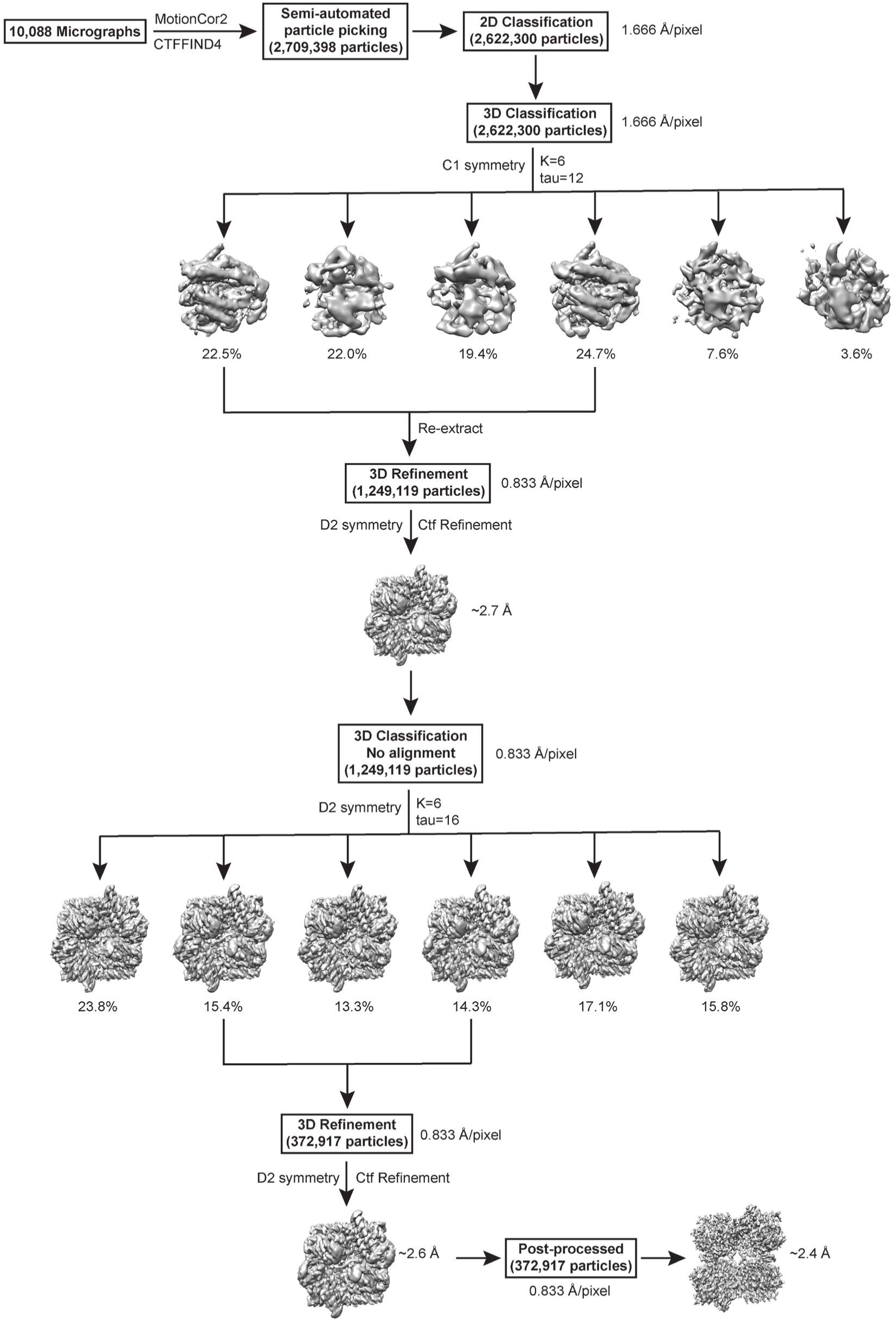
Flowchart for cryoEM data processing of NADH-bound COQ7:COQ9 complex. Details can be found in the image analysis and 3D reconstruction section of the Methods.

**Extended Data Figure 6.**
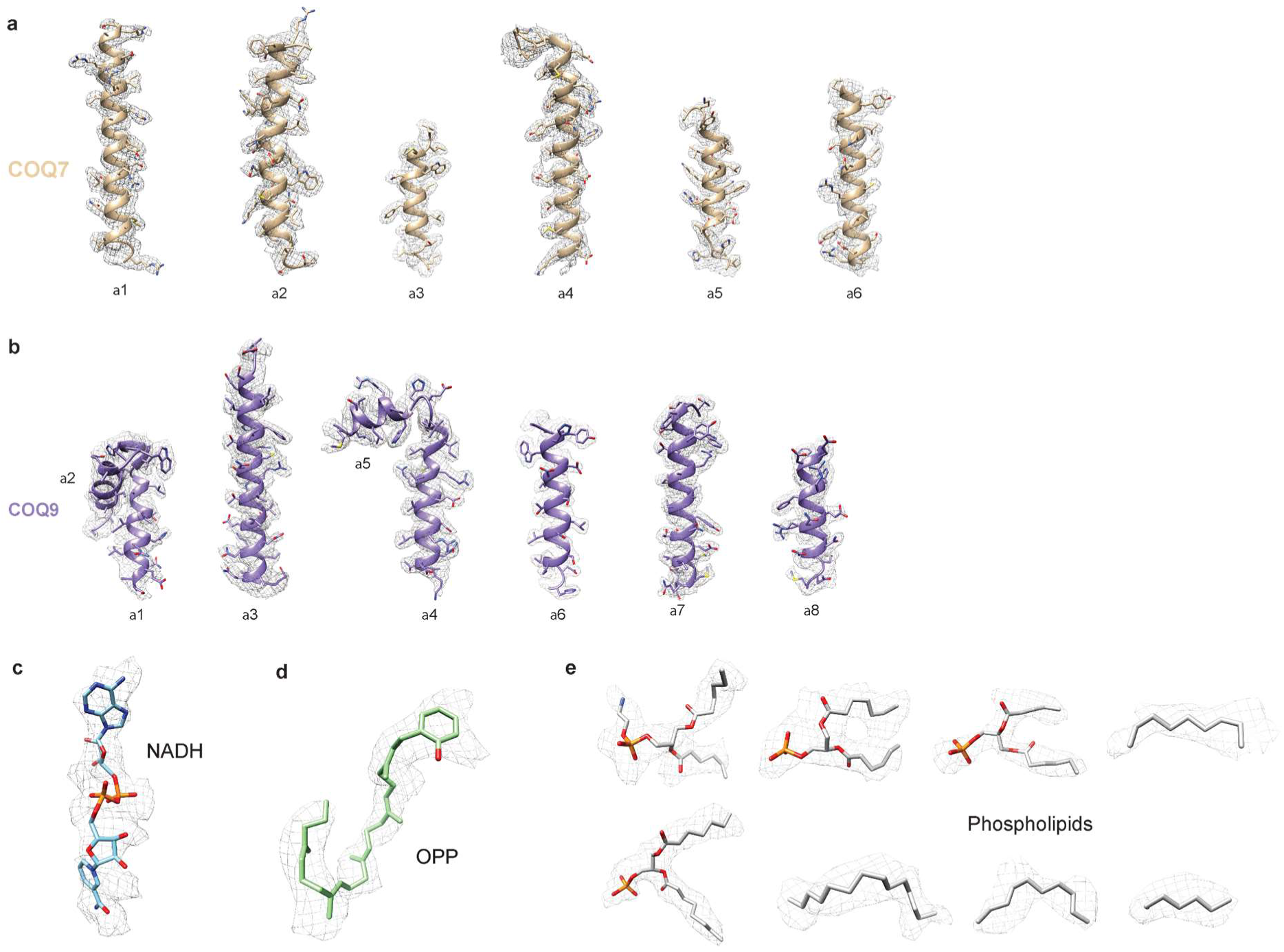
Snapshots of map versus model agreement by secondary structure. Refined atomic models fit well into corresponding densities. **a,b**, Representative densities include the ɑ-helical regions of COQ7 (beige) and COQ9 (purple) are shown in the context of the atomic model with side chains are shown as sticks and the backbone as ribbons. **c,d,e**, Representative regions of density from the electron density map for NADH (cyan), OPP (green), and phospholipid (gray) molecules are presented at the bottom panel.

**Extended Data Figure 7.**
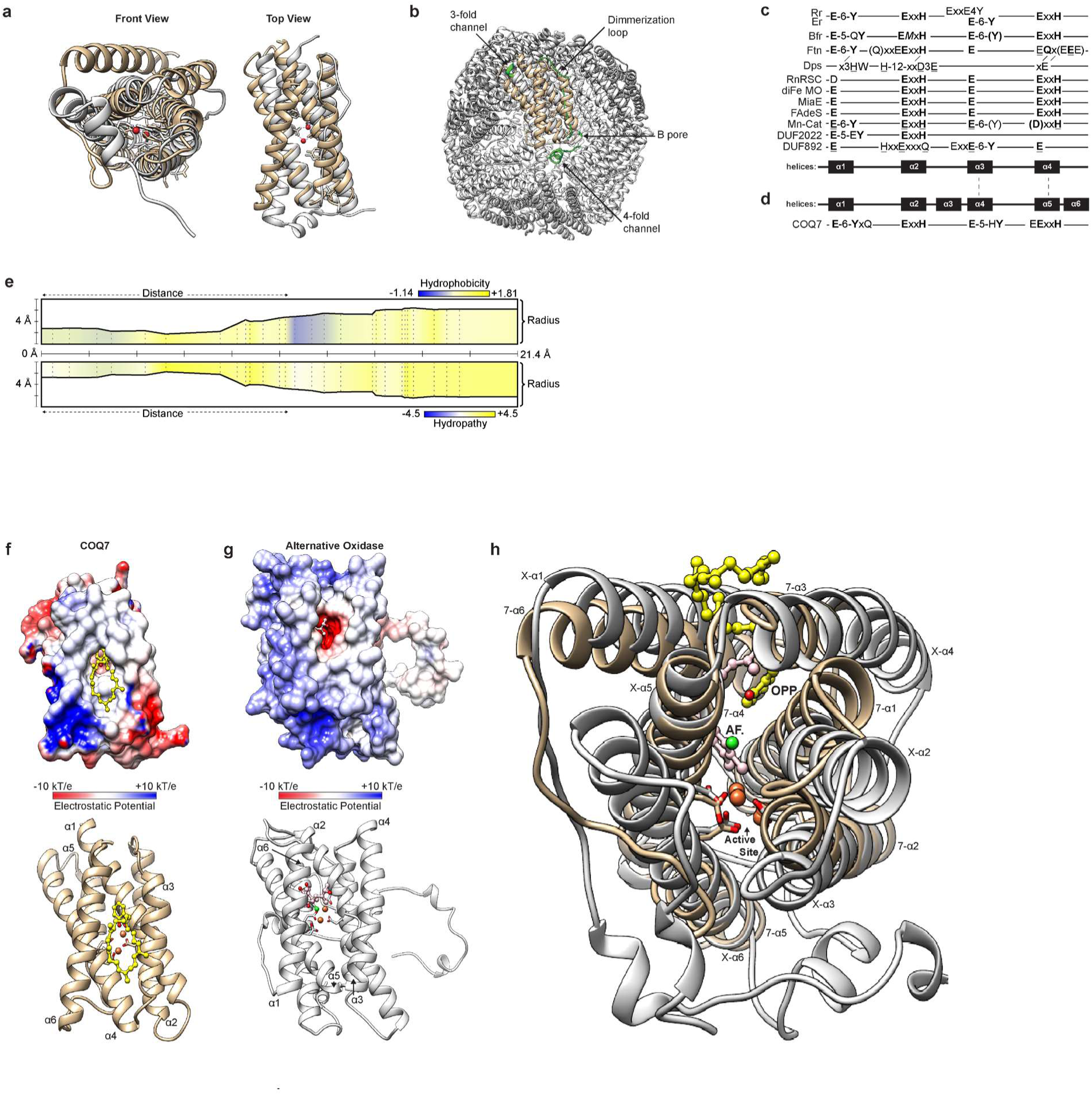
Comparison of COQ7 to bactoferritin and alternative oxidase. **a**, Structural comparison of cryoEM COQ7 (tan) to bactoferritin monomer (gray) PDB:4AM2 **b**, Structural features known to be important for oligomerization and function of ferritins but mutated in COQ7 are shown in green. **c**, Conservation of the (E-(X6)-Y-(X22)-E-(X2)-H-(X48)-E- (X6)-Y-(X28)-E-(X2)-H) motif among diiron proteins. **d**, topology of COQ7 in comparison to other diiron proteins. **e**, Properties of the hydrophobic channel leading to the COQ7’s active site calculated by MOLEonline. Panels c and d were reproduced with changes from Andrews, S.C^27^. **f**, Surface and ribbon view of COQ7. **g**, Surface and ribbon view of alternative oxidase PDB:3VVA (AOX). **h**, Structural comparison of COQ7 and AOX. Bound aromatic Ascofuranone-derivative inhibitor (pink, AF.) and resolved diiron active site (red spheres) compared to COQ7 (tan) with bound OPP (yellow). Helices of COQ7 (7-α) and AOX (X-α) are numbered.

**Extended Data Figure 8.**
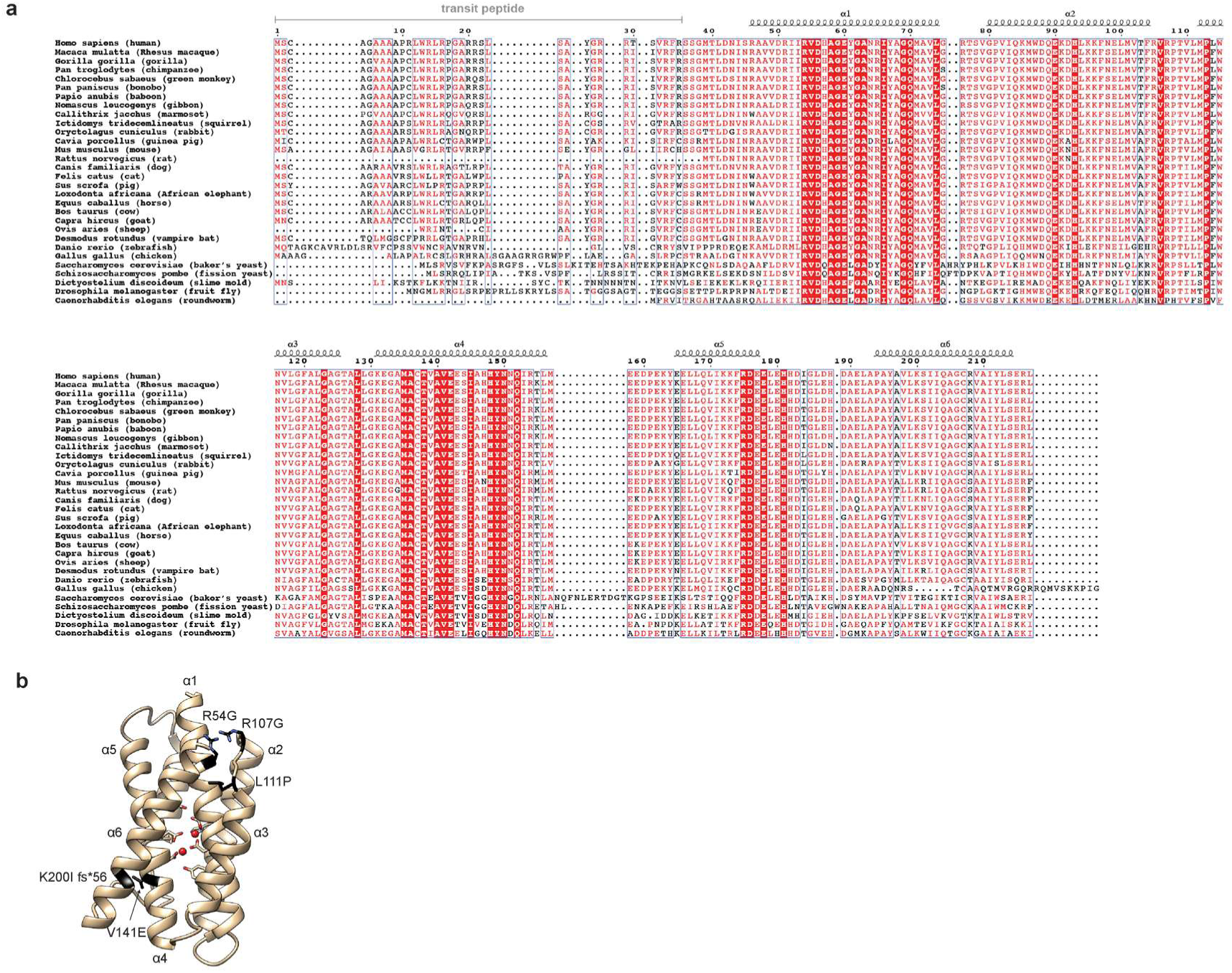
Evolutionary conservation of COQ7 and patient’s mutations found in COQ7. **a**, Multiple sequence alignment (MSA) showing the conservation of COQ7 residues. COQ7 sequences from Homo sapiens (human; Uniprot: Q99807), Macaca mulatta (rhesus macaque; Uniprot: A0A5F7ZCW4), Gorilla gorilla (gorilla; Uniprot: G3QJN6), Pan troglodytes (Chimpanzee; Uniprot: H2QAN4), Chlorocebus sabaeus (green monkey; Uniprot: A0A0D9R654), Pan paniscus (bonobo; Uniprot: A0A2R9BAP6), Papio anubis (baboon; Uniprot: A0A096N6T5), Nomascus leucogenys (gibbon; Uniprot: A0A2I3H0N0), Callithrix jacchus (marmoset; Uniprot: F6XXM7), Ictidomys tridecemlineatus (squirrel; Uniprot: I3M379), Oryctolagus cuniculus (rabbit; Uniprot: G1SGS8), Cavia porcellus (guinea pig; Uniprot: H0VHI5), Mus musculus (mouse; Uniprot: P97478), Rattus norvegicus (rat; Uniprot: Q63619), Canis familiaris (dog, Uniprot: E2RF61), Felis catus (cat; Uniprot: M3WJK9), Sus scrofa (pig; Uniprot: A0A287A0B5), Loxodonta africana (African elephant; Uniprot: G3STP5), Equus caballus (horse; Uniprot: F6YA13), Bos taurus (cow; Uniprot: Q2TBW2), Capra hircus (goat; Uniprot: A0A452EGI8), Ovis aries (sheep; Uniprot: W5PPJ7), Desmodus rotundus (Vampire bat; Uniprot: K9IHU0), Danio rerio (zebrafish; Uniprot: F1QW05), Gallus gallus (chicken; Uniprot: F1NIP2), Saccharomyces cerevisiae (baker’s yeast; Uniprot: P41735), Schizosaccharomyces pombe (fission yeast; Uniprot: O74826), Dictyostelium discoideum (slime mold; Uniprot: Q54VB3), Drosophila melanogaster (fruit fly; Uniprot: Q9W3W4), and Caenorhabditis elegans (roundworm; Uniprot: P48376) are aligned. Complete conservation of a given amino acid is indicated with red boxes. Secondary structural elements observed in the cryoEM structure of COQ7 are shown as coils for α -helices. **b**, Structure of COQ7 with detrimental mutations identified in patients based on ClinVar.

**Extended Data Figure 9.**
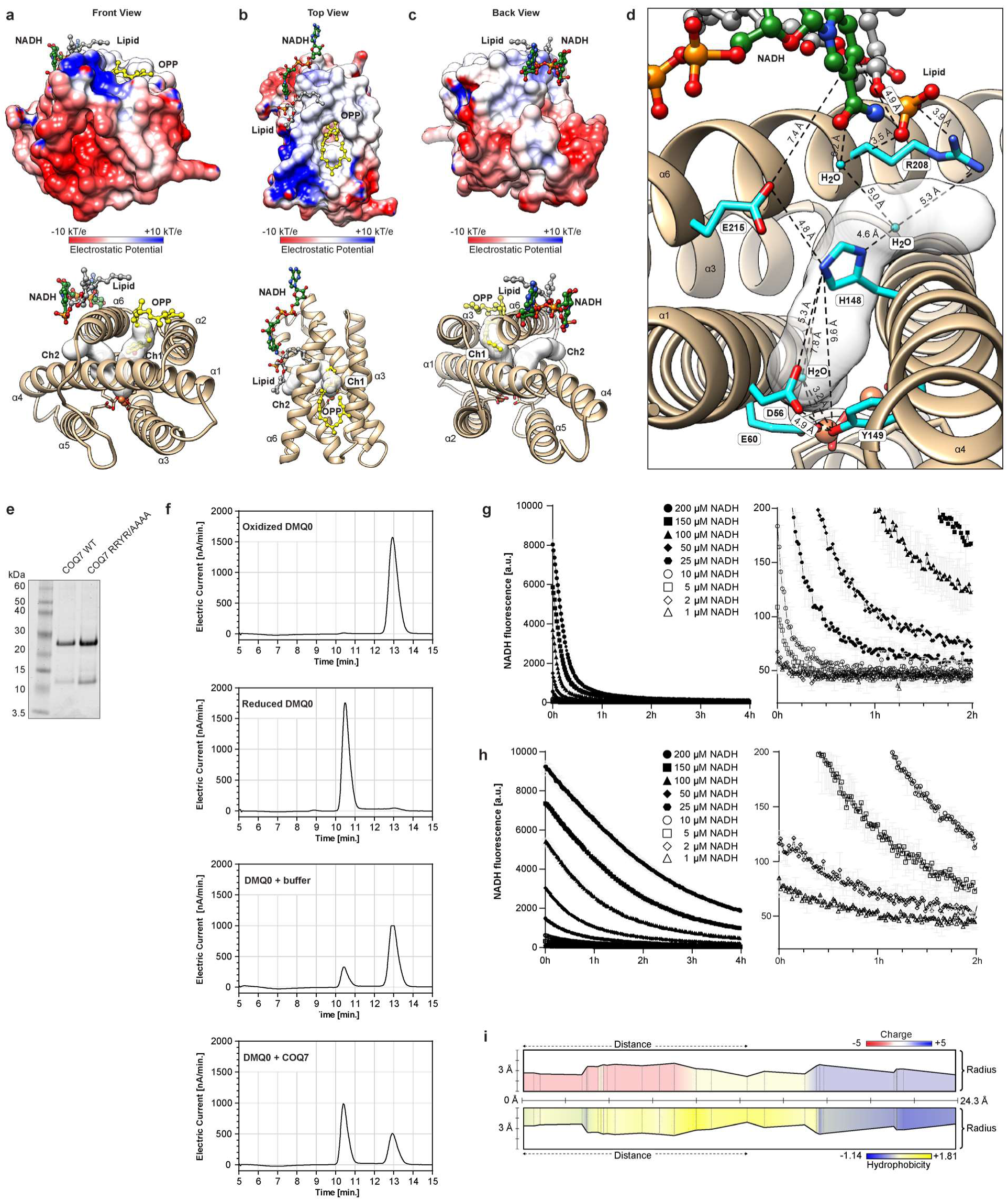
Additional details of NADH binding and utilization by COQ7. **a, b, c, d**, Additional poses presenting NADH binding site on COQ7 and channels leading to the active site. The OPP binding channel is labeled ch1 and the hypothesized charge transfer channel is labeled ch2. **e**, SDS-PAGE gel showing digitonin purified His6-GB1-^Nd38^COQ7 WT and His6- GB1-^Nd38^COQ7^R51A/R208A/Y212A/R216A^ mutants. **f**, HPLC-ECD analysis showing COQ7-dependent reduction of the quinone substrate DMQ0 by NADH present in the buffer. **g,h**, *In vitro* NADH titration experiments with COQ7 WT and COQ7 RRYR/AAAA (mean ± s.d., n = 3). Panels on the right are zooms in of the panels on the left. **i**, Properties of the channel **ch2** calculated by MOLEonline.

**Extended Data Figure 10.**
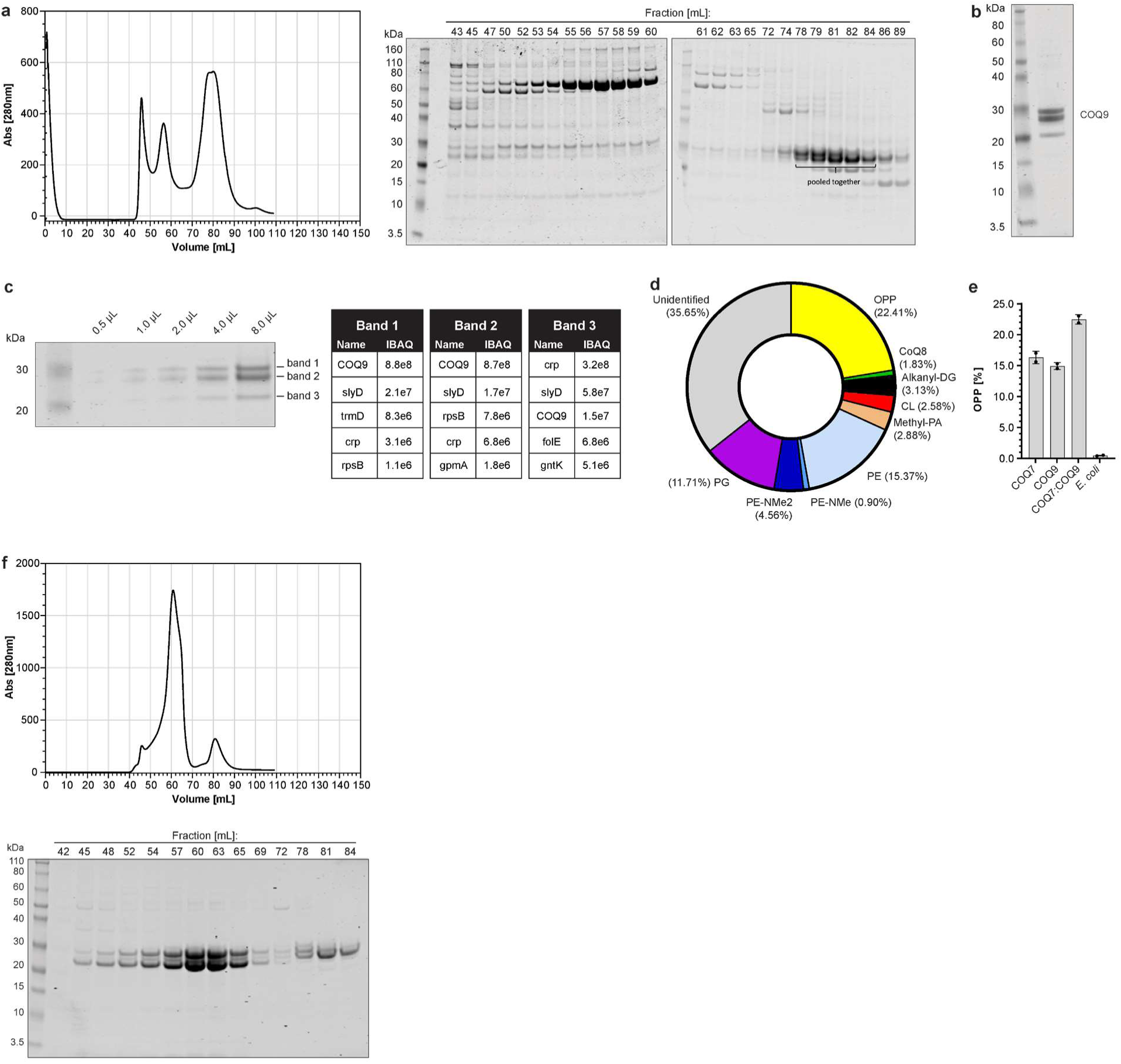
Analysis of COQ7:COQ9 dimer and tetramer. **a**, Size exclusion chromatography trace of His6-^Nd79^COQ9^W240K^ co-expressed with GB1- ^Nd38^COQ7. **b**, SDS-PAGE gel showing composition of the pooled fractions. **c**, LC-MS/MS identification of proteins in the bands. No COQ7 was detected. **d**, Lipids detected by LC-MS in the purified GB1-^Nd38^COQ7:His6-^Nd79^COQ9 complex. **e**, LC-MS measured level of OPP in individually purified His6-GB1-^Nd38^COQ7, His6-^Nd79^COQ9, GB1-^Nd38^COQ7:His6-^Nd79^COQ9 or *E.coli* cells. Data expressed as % of total lipid signal of each sample (mean ± s.d., n = 2).

**Extended Data Figure 11.**
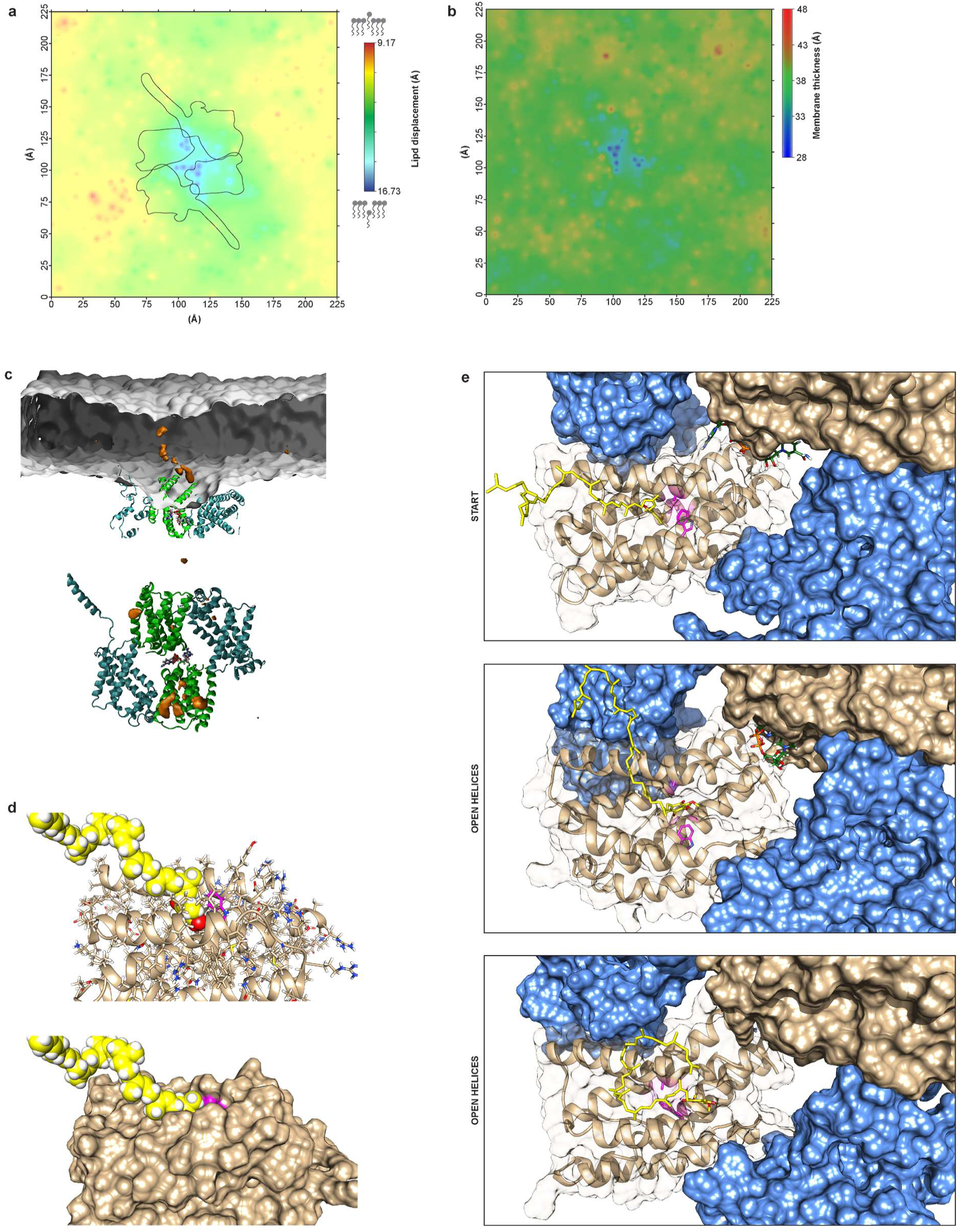
Additional data from COQ7:COQ9 MD simulation. **a**, Map of the deformation of the top membrane leaflet induced by the COQ7:COQ9 tetramer. **b**, Map of the changes of membrane thickness induced by the COQ7:COQ9 tetramer. **c**, Time-averaged density of CoQ10 molecules represented as brown volume/surface. **d**, Two views of a CoQ10 molecule modeled into COQ7 based on the OPP observed in the CryoEM structure. **e**, Additional representation of COQ7 conformational changes and CoQ10 movement visible in Fig. 5d.

**Extended Data Table 1.** CryoEM data collection, refinement, and validation statistics.

|  | NADH-bound COQ7:COQ9 complex<br>(EMD-25413)<br>(PDB 7SSS) | COQ7:COQ9 complex only<br>(EMD-25412)<br>(PDB 7SSP) |
| --- | --- | --- |
| Data collection and processing |  |  |
| Microscope | FEI Titan Krios | FEI Talos Arctica |
| Camera | Gatan K3 Summit | Gatan K3 Summit |
| Magnification | 105,000x | 36,000x |
| Voltage (kV) | 300 | 200 |
| Electron exposure (e<sup>-</sup>/Å<sup>2</sup>) | 65 | 58.8 |
| Defocus (μm) | -0.3 to -1.2 | -0.5 to -1.5 |
| Pixel size (Å) | 0.833 | 1.14 |
| Symmetry imposed | D2 | D2 |
| Micrographs (no.) | 10,088 | 1,395 |
| Initial particle images (no.) | 2,709,398 | 574,760 |
| Final particle images (no.) | 372,917 | 61,526 |
| Map resolution (Å) | 2.4 | 3.5 |
| FSC threshold | 0.143 | 0.143 |
| Map resolution range (Å) | 2.2 to 6.7 | 3.4 to 5.9 |
| Refinement |  |  |
| Initial model used (PDB code) | COQ7 (*de novo*) – COQ9 (6AWL) | COQ7 (*de novo*) – COQ9 (6AWL) |
| Model resolution (Å) | 2.4 | 3.5 |
| FSC threshold | 0.143 | 0.143 |
| Map sharpening *B* factor (Å<sup>2</sup>) | -95.99 | -156.07 |
| Model composition |  |  |
| Nonhydrogen atoms | 12,190 | 10,930 |
| Protein residues | 1,416 | 1,376 |
| Ligands | 4 |  |
| Lipids | 32 |  |
| Cofactor | 2 |  |
| Water | 170 |  |
| B factors (Å<sup>2</sup>) |  |  |
| Protein | 25.11 | 22.16 |
| Ligands | 43.18 |  |
| Water | 30.84 |  |
| R.m.s. deviations |  |  |
| Bond lengths (Å) | 0.004 | 0.004 |
| Bond angles (Å) | 0.449 | 0.611 |
| Validation |  |  |
| MolProbity score | 1.27 | 1.67 |
| Clashscore | 4.01 | 4.52 |
| Poor rotamers (%) | 0.0 | 0.0 |
| Ramachandran plot |  |  |
| Favored (%) | 97.63 | 93.15 |
| Allowed (%) | 2.37 | 6.85 |
| Disallowed (%) | 0.0 | 0.0 |

